# Genetic diversity and variation in heterozygosity in Macadamia

**DOI:** 10.1101/2025.09.22.677927

**Authors:** Sachini Lakmini Manatunga, Agnelo Furtado, Bruce Topp, Mobashwer Alam, Patrick J. Mason, Robert J Henry

## Abstract

**Background:** Macadamia is a diploid (2n =28) rainforest tree of the Proteaceae family, native to Australia. Single-nucleotide polymorphisms (SNPs) have proven to be effective tools for evaluating genetic diversity and heterozygosity. SNPs from the whole genome were used to determine allelic diversity within different species and breeding populations, and to investigate the heterozygosity of genotypes

**Results:** A total of 349 samples, including wild and domesticated genotypes, were whole genome sequenced, generating 4,180,786 – 7,661,269, 2,446,762 - 7,356,187, 2,829,944 - 4,391,600, 1601519 - 3,182,089 and 3,123,715 to 9,620,640 single nucleotide polymorphism (SNP) positions at the whole genome level for *M. integrifolia*, *M. tetraphylla*, *M. ternifolia*, *M. jansenii* and the domesticated cultivars and breeding selections respectively. The overall SNP diversity for wild *M. integrifolia*, *M. tetraphylla*, *M. ternifolia* and *M. jansenii* was 4.43%, 4.33%, 1.80% and 1.1%, respectively. Domesticated *M. integrifolia* showed an overall SNP diversity of 3.4%, 1.03% lower than wild *M. integrifolia*, providing evidence of a genetic bottleneck during domestication. Genetic diversity at the individual level for the domesticated gene pool varied from 0.4% to 1.24%, with an overall diversity of 6.10%. Results across different breeding groups revealed that the Australian breeding selections (NMBPA) had the highest diversity, and HAES cultivars had the lowest diversity. Wild *M. integrifolia* and *M. tetraphylla* exhibited heterozygosity ranging from 0.36% to 0.71% and 0.30% to 0.71% respectively. The two non-edible species, *M. ternifolia* and *M. jansenii*, displayed relatively low heterozygosity (From 0.29% to 0.42% and 0.19% to 0.34% respectively). Domesticated macadamia exhibited a wide range of heterozygosity (0.33% to 1.15%). Additionally, heterozygosity correlated with yield efficiency (kernel yield/tree volume), cumulative kernel yield at age 8 and cumulative nut in shell (NIS) yield at age 8 and showed that performance increased with heterozygosity to an optimal level and declined thereafter.

**Conclusion:** These findings confirmed that there is substantial genetic variation among the different macadamia genotypes. The results also revealed that the greatest diversity is present in wild germplasm. The data will greatly support future genome-wide studies and breeding programs.

## Background

Genetic diversity, the variation in genetic information within a population or species, is the basis of biodiversity and evolution [1–4]. The generation and maintenance of genetic diversity occurs mainly through recombination in sexual reproduction, which leads to higher genetic variation in a population than in plants propagated by an asexual/vegetative method [4, 5]. In plants, genetic diversity is determined by several factors such as natural selection, migration, mutation and genetic drift [6]. A population with low genetic diversity faces an increased risk of population decline or extinction compared to a population with rich genetic diversity because it lacks the ability to adapt effectively to environmental changes such as climate change, pests, diseases, and habitat alteration [4, 7, 8]. Therefore, maintaining genetic diversity within a population or species can enhance its long-term survival and evolutionary adaptability [5].

Macadamia is a recently domesticated nut crop of the Proteaceae and subfamily Grevilleoideae. The genus *Macadamia* contains four species. *M. integrifolia* has the widest natural distribution in Australia, followed by *M. tetraphylla* and *M. ternifolia*, which are spread mainly in New South Wales and north of Brisbane, respectively. *M. jansenii* is the smallest isolated population found only in Bulburin National Park, north of Bundaberg [9]. *M. integrifolia* and *M. tetraphylla* are edible species used in commercial cultivar development [10, 11]. The other two species are inedible due to the presence of cyanogenic glycosides [11, 12]. Macadamia is partially self-incompatible, which means the species mostly exhibits self-compatibility but not entirely [13]. This mechanism prevents self-fertilisation and promotes cross-pollination to increase diversity.

The study of genetic diversity has depended on sequencing technology advancements, enabling researchers to obtain high-quality genomic data from different plant species [4]. Sequencing technology allows the detection of millions of genome-wide DNA polymorphisms such as single-nucleotide polymorphisms (SNPs), multi-nucleotide polymorphisms (MNPs), insertions and deletions (Indels) [14]. Among these, SNPs are the most abundant and robust variations in the plant genome [15]. They are in the form of either ‘heterozygous SNPs’ or ‘homozygous SNPs’ and are invaluable resources for genomic approaches and crop improvement [16].

Many genetic diversity studies based on SNPs identified from whole-genome sequencing data have been reported for crops such as bitter gourd [17], tobacco [15], cucumber [18], and plum [19]. SNPs were identified using SNP calling tools such as Genome Analysis Toolkit (GATK) [20], SAMTOOLS [21], SOAP2 [22] and a basic variant detection algorithm in the CLC software. For macadamia, genetic diversity has been assessed extensively by using different molecular markers [23–29]. Few studies have investigated diversity based on SNPs identified from whole-genome sequence data. The genetic diversity of the *M. jansenii* population was assessed using SNPs identified from a basic variant detection tool from whole genome sequencing data, which revealed low diversity within the species [30]. A study by Lin *et al.,* reported the diversity for 112 accessions, including wild and cultivars and selected lines based on SNPs [31]. Furthermore, a recent study reported the genetic diversity of Chinese macadamia germplasm [32]. However, to our knowledge, no studies have investigated population-scale allelic diversity in macadamia.

Heterozygosity is a common way of measuring genetic resilience [33]. The knowledge of heterozygosity in different species and populations is crucial for conservation biology and also to enhance agricultural productivity and long-term sustainability [34]. In general, a population with higher heterozygosity has greater adaptability against biotic and abiotic stress than a population with low heterozygosity [34]. Recently, attention has been geared toward the use of genome-wide heterozygosity in plants. In macadamia, initially, isozyme marker data were used to determine heterozygosity in all species except *M. jansenii* [35]. Since then, many studies have reported the heterozygosity for both wild and domesticated cultivars [23, 26, 28, 29, 32, 36–41]. Molecular marker analyses have indicated a high level of heterozygosity in macadamia [40]. However, the variation in heterozygosity of individual plants in wild and domesticated macadamia remains largely unknown.

The objective of this present study was to better understand allelic diversity within the four macadamia species and in domesticated material and to compare the diversity among the individuals in wild and domesticated samples. Another objective was to investigate heterozygosity based on the SNPs identified by whole-genome sequencing and to understand the relationship between heterozygosity and phenotypic traits important in crop production.

## Methods

### Plant materials

A total of 349 macadamia samples were analysed in this study. They included 150 wild macadamia accessions of all four species [*M. integrifolia* (n = 48), *M. tetraphylla* (n = 56), *M. ternifolia* (n = 23), and *M. jansenii* (n = 23)] (Table S.1) and 199 macadamia cultivars from different breeding programs (Table S. 2) and breeding selections from an Australian macadamia breeding program. Among wild material, 161 were collected from Maroochy Research Facility in Nambour and Tiaro in Queensland and Alstonville in NSW, Australia [42]. The remaining five samples were collected from a private collection at Limpinwood, NSW, Australia. All the cultivars and breeding selections were collected from ex-situ germplasm centres at Alstonville, Beerwah, Bundaberg, Nambour and Tiaro. Fully expanded young fresh leaves were collected in perforated bags and immediately placed under dry ice until they were stored in a −80 °C freezer at The University of Queensland, Australia.

### DNA extraction and sequencing

Genomic DNA was extracted from leaf tissues that were coarsely pulverised and then finely pulverised [pulverised under cryogenic conditions using a Qiagen tissue lyser (MM400, Retsch, Germany)] using a modified version of the cetyltrimethylammonium bromide (CTAB) method [43]. The quality of the DNA samples was evaluated using a Nanodrop spectrophotometer (Nanodrop Technologies, Wilmington, DE, USA) by recording the absorbance ratios at 260/280 and 260/230. The quantity of the DNA samples was assessed by running a 0.7% agarose gel with SYBR Safe staining (Thermo Fisher Scientific). Whole genome short read sequencing was performed by BGI Hong Kong using the advanced DNBSEQ-G400 platform from MGI (MGI Tech Co., Ltd, Shenzhen, China). A high-quality PCR-free library was prepared, and sequencing was performed to produce 150 bp paired-end reads.

### Data processing

Raw paired-end reads were imported to CLC Genomics Workbench 23.0.05 (CLC-GWB, CLC-Bio, QIAGEN). Reads of all the samples were trimmed using a quality limit of 0.01 with default parameters. Most of the resulting trimmed reads (More than 98%) had a Phred score >25.

### The variant analysis

All data analysis was undertaken using CLC Genomics Workbench 23.0.05 (CLC-GWB, CLC-Bio, QIAGEN). For variant analysis, reference genomes were selected for each of the four species. The *M. integrifolia* (GWHESFF00000000) reference genome was selected for all the wild (n=48) and domesticated (n=39) *M. integrifolia* and for all the cultivars and breeding selections (n=199). Wild *M. tetraphylla* (n=56), *M. ternifolia* (n=23) and *M. jansenii* (n=23) were analysed using their respective reference genomes: *M. tetraphylla* (GWHESFG00000000), *M. ternifolia* (GWHESEN00000000) and *M. jansenii* (GWHESFI00000000) [44]. Trimmed paired end reads of all the samples were individually mapped against their respective reference genomes using the “Map to reference” tool. Mapping titration was performed at four different stringency levels of read length fraction (LF) and read similarity fractions (SF) (LF 1.0 SF 0.95, LF 1.0 SF 0.9, LF 0.9 SF 0.9, LF 0.8 SF 0.8) for the selected three genotypes from each species separately to select optimal setting for the read mapping (Figure S. 1). LF 1.0 and SF 0.9 were selected as the optimal mapping settings for all the read alignments based on the mapping percentage. The parameter for read mapping as follows: masking mode “No masking”, match score “1”, mismatch cost “2”, insertion cost “3”, deletion cost “3”, non-specific match handling “map randomly”, execution mode “standard” and minimum seed length “15”. PCR derived duplicate reads generated during sequencing library preparation were removed from the read alignments to prevent capture of false-positive SNPs in subsequent variant analysis. Then, broken paired-end reads were excluded in read mapping and “InDels and structural variants” analysis was performed to identify erroneous structural variants such as insertions, deletions, inversions and tandem duplications. Local realignment was performed to improve the read mapping by re-mapping reads around insertions and deletions. This process corrects the misalignment by using information from nearby reads. Then, read mapping was used to perform “Fixed ploidy variant detection” to identify SNP variants in comparison to the respective reference genome. Frequency titration was performed to select the optimal setting for variant calling (Figure S. 2) at five different levels of minimum frequency percentage (40%, 35%, 30%, 25% and 20%). A minimum frequency percentage of 25 was selected as the optimal setting based on SNP variant positions. The parameters of fixed ploidy analysis are as follows: Required variant probability “90.0”, Minimum coverage “10”, Minimum count “3”, Base quality filter “Yes”, Neighbourhood radius “5”, Minimum central quality “20”, Minimum neighbourhood quality “15”, and Direction frequency (%) “5.0”.

Following this, MNV, insertion, deletion, replacement, heterozygous SNPs ranged between 25%-75%, homozygous SNPs at 100% frequency with more than 20X coverage were filtered for whole genome and CDS region individually for all the samples.

### Diversity analysis

Diversity analysis was conducted separately for 16 different populations, namely, wild *M. integrifolia* (n=48), wild *M. tetraphylla* (n=56), *M. ternifolia* (n=23), *M. jansenii* (n=23), domesticated *M. integrifolia* (n=39), cultivars from Hawaiian Agricultural Experiment Station (HAES) (n=37), cultivars from Hidden Valley Plantation (HVP) (n=12), cultivars from Heritage (n=17), cultivars by Norm Greber (n=9), Cultivars by Backer (n=4), cultivars by Malawi (n=2), cultivars by Ian McConachie (n=7), Australian national breeding program accessions (NMBPA) (n=108), whole domesticated gene pool (n=199) and cultivars and breeding selections grouped in clade I (n=4) and clade II (n=195) in nuclear phylogenetic analysis performed in our previous publication.

All data analysis was undertaken using CLC Genomics Workbench 23.0.05 (CLC-GWB, CLC-Bio, QIAGEN). Diversity analysis was performed based on heterozygous SNPs, ranging between 25%-75%, and homozygous SNPs at 100% frequency, with more than 20X coverage in whole. For the individual genotype of the selected population, heterozygous SNPs and homozygous SNPs at 100% frequency with more than 20X coverage were merged into a single file. For the specific population, all the individual merge files of all the genotypes are combined using “Frequency threshold (%) = 0” (Figure S. 3). Non-diverse homozygous SNPs were separated using the filters “Sample count with number of total samples” and “Zygosity contains Homozygous”. Non-diverse heterozygous SNPs were separated using the filters “Sample count with number of total samples” and “Zygosity contains Heterozygous”. Non-diverse homozygous SNPs and heterozygous SNPs were merged and excluded to keep only the diverse SNPs for the analysis.

Definitely diverse SNPs (polymorphic positions where genotypes include either a mix of heterozygous and homozygous positions, or a combination of heterozygous, homozygous positions, and the other genotypes have the reference allele) were further filtered and separated from the whole set of diverse SNPs. The remaining diverse SNPs, which are known as “Doubtful diverse SNPs” (diverse positions where genotypes are either all heterozygous or some genotypes have heterozygous positions while the rest carry reference allele and homozygous diverse positions), were filtered for homozygous and heterozygous SNP positions separately. Data noise resulting from unmapped reads and the failure of variant calling parameters was identified in doubtful diverse SNP positions by running a mutation analysis pipeline [45]. Then, the heterozygous SNP positions that were similar to the reference in all the genotypes were combined and filtered against total diverse SNPs using the filter option “Keep variants with exact match found in the track of known variants”. Homozygous SNP positions that were similar to the reference were combined and filtered against total diverse SNPs using the filter option “Keep variants with exact match found in the track of known variants”. Total diverse heterozygous SNP positions were calculated by adding the number of heterozygous SNP positions similar to the reference in all the genotypes and definite diverse SNPs. Total diverse SNP positions were calculated by adding the number of heterozygous SNP positions that are similar to the reference in all the genotypes, homozygous SNP positions that were similar to the reference and definite diverse SNPs. Allelic diversity was calculated as a percentage of polymorphic loci to the total genomic size. Individual diversity values were calculated by filtering “Origin track with the name of the genotype”. Accession-specific, unique, polymorphic heterozygous SNP positions were calculated by filtering total diverse heterozygous SNPs with “Sample count <= 1”. Accession-specific, unique, polymorphic homozygous SNP positions were calculated by filtering total diverse homozygous SNPs with “Sample count <= 1”.

### Heterozygosity calculation

The heterozygosity for each sample was calculated as a percentage of heterozygous SNV positions in the reference genome. Heterozygous SNV positions were identified by filtering ‘Reference allele contains No’.

### Phenotypic data collection standardization and correlation analysis

Historical phenotypic datasets were utilized in this study. This study used data for six traits (Cumulative nut in shell (NIS) yield at age 8, cumulative kernel yield at age 8, kernel recovery (kernel weight/nut in shell weight), canopy volume at age 6, tree height at age 6 and yield efficiency (kernel yield /tree vol)) across 79 genotypes (Table S. 3) comprising both parents and the second generation progeny from the Australian breeding program, planted across 14 trials at nine locations [46]. Phenotypic data were normalised to control for location and analysed using ASReml-R as described in Topp *et al.,* 2014 [47]. Pearson’s correlation was used to assess the relationship between heterozygosity and the six measured traits.

## Results

### Sequencing and mapping of reads to the reference genome

The sequencing of 349 macadamia accessions generated raw reads between 98,50,980 – 260,592,896 with a sequence coverage range between 18.23X - 50.11X (Table S. 4). Trimmed reads at 0.01 quality limit generated reads between 93,695,956 - 247,617,841.

The mapping was performed with four reference genomes. For wild *M. integrifolia* populations (n=48), wild macadamia trees from mixed/hybrid population (n=4), wild macadamia trees of uncertain species (n=9), a wild hybrid (n=1), a wild admixture (n=1) and for all cultivars and breeding selection (n=199), the mapping of trimmed paired-end reads from each of the accessions on to the *M. integrifolia* genome varied from 70.2% to 93.9% (Table S. 5). The lowest mapping percentage was recorded for Mac_318, which is known to be a wild macadamia tree of uncertain species. For the wild *M. tetraphylla* population (n=56), the mapping of trimmed paired end reads from each accession to the *M. tetraphylla* genome varied from 80.8% to 93.4% (Table S. 6). The mapping of trimmed paired end reads from each *M. ternifolia* accession to the *M. ternifolia* genome varied from 76.5% to 95.1% (Table S. 7). For the *M. jansenii* population (n=23), the mapping of trimmed paired end reads from each accession to the *M. jansenii* genome varied from 87.3% to 95.4% (Table S. 8).

### Detection of DNA polymorphisms

Among the wild *M. integrifolia* population, there were 4,180,786 – 7,661,269 SNP positions across 48 genotypes, revealing 5.39 – 9.88 SNP positions per one kb of *M. integrifolia* genome (Figure S. 4). The total number of MNPs, insertions, deletions and replacements were in the range between 221,995 to 326,706, 355,810 to 472,711, 366,919 to 536,547 and 31,772 to 43,398, respectively (Table S. 9). The highest total number of SNPs was recorded for Mac_035, and the least was recorded for Mac_171. Mac_091 recorded the highest heterozygous SNPs, whereas Mac_035 recorded the highest 100% homozygous SNPs (Table S. 10). The lowest number of heterozygous SNPs was recorded for Mac_250 and the lowest homozygous SNPs for Mac_130. At the CDS level, the total number of SNP positions was in the range between 159,772 to 273,486 (Table S. 9).

For the wild *M. tetraphylla* population, there were 2.9M – 8.8M variant positions in relation to the reference genome for the 56 genotypes. At the whole genome level, SNPs (2,446,762 to 7,356,187) (3.07 SNP/kb – 9.25 SNP/kb) were the major type of variants, followed by deletions (203,140 to 574,046), insertions (169,567 to 477,646), MNPs (110,618 to 395,403) and replacements (12,586 to 48,490) (Figure S.5, Table S. 11). The highest number of SNPs (12,550,003) was recorded for Mac_236 and the least recorded for MacP_15 (4,832,434) (Table S. 12) compared to the reference *M. tetraphylla* genome. Heterozygous SNPs ranged from 4,742,286 – 11,333,633 with the highest recorded for Mac_236. Homozygous SNPs (100%) ranged from 90,148 – 2,777,121 with the highest recorded for Mac_030, which is believed to be a planted tree from wild trees. Within the coding sequences, the total number of SNP positions were between 83,793 and 277,573(Table S. 11).

Within the 23 samples of the *M. ternifolia* population, 3.6M - 5.2M variant positions were identified in the whole genome. There were 3.62 – 5.62 SNP positions per one kb of *M. ternifolia* genome (Figure S. 6). The total number of MNPs, insertions, deletions and replacements ranged from 88,709 to 170,484; 173,145 to 335,609; 200,108 to 375,617; 12,833 to 26,513, respectively (Table S. 13). Mac_300 recorded the highest heterozygous SNPs (6,556,435), and Mac_126 recorded the lowest heterozygous SNPs (4,649,136) (Table S. 14). Mac_332 recorded the highest homozygous SNPs (1,411,730), and MacP_12 recorded the lowest homozygous SNPs (11). At the CDS level, between 1.1M – 1.8M variant positions were identified, with SNPs representing the predominant variant type ranging from 106,034 to 166,120 (Table S. 13).

Among the 23 *M. jansenii* genotypes the total number of variant positions ranged from 1.9M to 3.8M. The density of SNPs across the *M. jansenii* genome ranged from 2.07 to 4.12 per kilobase (Figure S. 7). MNP positions varied from 66,284 to 119,204. Insertions were from 121,641 to 241,370, while deletions ranged from 144,258 to 281,908. Replacements were comparatively few, ranging between 8,330 and 18,418 (Table S. 15). The highest total number of SNPs was recorded for Mac_282, and the least was recorded for MacP_16. Mac_282 recorded the highest heterozygous SNPs, whereas MacP_02 recorded the highest 100% homozygous SNPs (Table S. 16). Within the coding sequences, the total number of SNP positions were between 59, 297 and 111,702 (Table S. 15).

Variant analysis across domesticated macadamia genotypes revealed a wide range of polymorphism. The total number of variant positions identified ranged from 3.8M to 11.3M. Domesticated cultivars and breeding selections exhibited SNP densities ranging from 4.02 to 12.4 SNPs per one kb of *M. integrifolia* genome (Figure S. 8). MNPs ranged from 97,087 to 476,580, insertions ranged from239,499 to 537,367, deletions ranged from 274,691 to 613,841 and replacements ranged from 18,198 to 55,693 (Table S. 17). Mac_333 from the Australian National Macadamia Breeding Program Accession (NMBPA), known as the wild domestic hybrid, recorded the highest number of SNPs and heterozygous SNPs (Table S. 18). The cultivar known as ‘Minimacca’ (Mac_347) recorded the highest number of homozygous SNPs (100%), whereas Mac_135, ‘HAES 741’ recorded the lowest number of homozygous SNPs. The total number of SNV variant positions in CDS regions were in the range between 107,053 and 366,556 (Table S. 17).

### Diversity analysis and comparison

Diversity analysis was performed based on the percentage of polymorphic loci for different groups of populations. Allelic diversity for the wild *M. integrifolia* population was presented in Table S. 19. The result showed that the overall diversity within the population was 4.43% with individual values ranging from 0.53% to 0.97% (Figure S. 9), indicating 0.53% to 0.97% of the genome consists of polymorphic loci. Mac_163 exhibited the fewest polymorphic positions compared to the reference genome. The highest diversity was observed for Mac_035. The highest accession-specific, unique, polymorphic heterozygous SNP positions (733,189) were found in Mac_344 (Table S. 20). The highest accession-specific, unique, polymorphic homozygous SNP positions (34,686) were found in Mac_250.

For the domesticated *M. integrifolia* population, allelic diversity ranged from 0.39% to 0.96%. Overall diversity was 3.40% for the 39 genotypes (Table S. 21) (Figure S. 10). The lowest and the second lowest were recorded for the cultivar HAES 741 (Mac_135 and Mac_195, respectively) with respect to its HAES 741-reference genome (*M. integrifolia*). The highest diversity was recorded for cultivar ‘Daddow’ (Mac_072). Accession-specific, unique, polymorphic heterozygous SNP positions ranged from 15,303 (Mac_055) to 1,497,179 (Mac_156) (Table S. 22). The highest and lowest number of accession-specific, unique, polymorphic heterozygous SNP positions recorded for two Hawaiian cultivars: HAES 800 (Mac_156) and HAES 807 (Mac_055), respectively. In addition, the highest number of accession-specific, unique, polymorphic homozygous SNP positions were found in cultivar D1 (Mac_087).

A comparison of allelic diversity between wild and domesticated *M. integrifolia* revealed greater diversity for wild germplasm. An independent two sample t test indicated a statistically significant difference between the two datasets (p value - 9.81367E-07) (Figure S. 11).

Wild *M. tetraphylla* populations showed an overall diversity of 4.33% across 56 genotypes. Individual diversity values indicated that 0.31% to 0.92% of the *M. tetraphylla* genome contained diverse positions within the population (Table S. 23) (Figure S. 12). MacP_015 recorded the lowest diversity, while Mac_030, a planted tree, recorded the highest. The number of accession-specific, unique, polymorphic heterozygous SNP positions ranged from 25,308 to 2,863,676 (Table S. 24). Accession-specific, unique, polymorphic homozygous SNP positions ranged from 123 to 614,911.

The *M. ternifolia* population recorded 0.36% to 0.55% diversity for 23 genotypes (Figure S. 13). Overall diversity was 1.80% (Table S. 25). The lowest diversity was observed in two genotypes: MacP_12, which is the reference genome accession, and Mac_126. In contrast, the highest diversity found in two genotypes: Mac_300 and Mac_310. The number of accession-specific, unique, polymorphic heterozygous and homozygous SNP positions ranged from 27,588 to 419,584 and zero to 15,430, respectively (Table S. 26).

For the isolated population of *M. jansenii*, overall diversity was 1.1%, with individual diversity ranging from 0.2% to 0.4% (Table S. 27) (Figure S. 14). The highest accession-specific, unique, polymorphic heterozygous SNP positions (94,407) were observed in MacP_04 (Table S. 28), while the highest accession-specific, unique, polymorphic homozygous SNP positions (2,103) were found in Mac_218.

Diversity analyses for domesticated cultivars and breeding selections showed that the overall diversity was 6.10% across 199 genotypes (Table S. 29). Genomic diversity at the individual level varied from 0.4% to 1.24% (Figure S. 15). The highest diversity was observed in Mac_348, which is a hybrid of wild x domestic, and the lowest diversity was observed in Mac_135, which is the reference genome accession. The number of accession-specific, unique, polymorphic heterozygous SNP positions ranged from 3,177 to 434,165 (Table S. 30). The number of accession-specific, unique, polymorphic homozygous SNP positions ranged from zero to 29,551. The results showed that 19 genotypes (Mac_005, Mac_016, Mac_093, Mac_102, Mac_121, Mac_133, Mac_135, Mac_137, Mac_160, Mac_162, Mac_166, Mac_176, Mac_186, Mac_190, Mac_195, Mac_203, Mac_269, Mac_285 and Mac_322) don’t have any accession specific unique, polymorphic homozygous SNP positions compared to the reference HAES 741 *M. integrifolia* genome.

We additionally analysed genetic diversity across different subgroups from the domesticated gene pool defined by their breeding program. Genetic diversity within the Hawaiian cultivars (n=37) ranged from 0.39% to 0.96%, with an overall diversity of 3.33% (Table S. 31) (Figure S. 16). The highest individual diversity was observed in cultivar HAES 853. The minimum number of accession-specific, unique, polymorphic SNP positions (Both heterozygous and homozygous) was recorded in HAES 849 with a total of 15,673 unique SNPs (Table S. 32). In the HVP cultivars (n=12), overall diversity was 2.31% (Table S. 33). Individual diversity ranged from 0.56% in Mac_295 (Cultivar A9/9) to 1.02% in Mac_061 (Cultivar A16) (Figure S. 17). Accession-specific, unique SNPs presented in Table S. 34. For heritage cultivars (n=17), individual diversity values varied with the lowest observed in Mac_151 and Mac_154 at 0.52% and the highest in Mac_230 at 1.05% (Table S. 35) (Figure S. 18). The cultivar ‘Beaumont’ recorded the highest number of accession-specific, unique SNPs (Table S. 36). Overall genetic diversity for cultivars bred by Norm Greber (n=9), Backer (n=4), Malawi (n=2), Ian McConachie (n=7) were 2.53% (Table S. 37) (Figure S. 19), 0.96% (Table S. 38) (Figure S. 20), 0.46% (Table S. 39) (Figure S. 21) and 2.03% (Table S. 40) (Figure S. 22) respectively. The highest number of accession-specific, unique, polymorphic SNP positions for cultivars bred by Norm Greber, Backer, Malawi, Ian McConachie were observed in Mac_187 (1,709,907) (Table S. 41), Mac_170 (1,979,851) (Table S. 42), Mac_158 (1,866,075) (Table S. 43) and Mac_070 (2,233,011) (Table S. 44). Australian breeding selections (NMBPA) showed 5.16% diversity across 108 genotypes, with individual values ranging from 0.43% to 1.23% (Table S. 45) (Figure S. 23). The maximum number of accession-specific, unique, polymorphic SNP positions observed in Mac_348 (Table S. 46). Allelic diversity across different breeding groups, based on the individual diversity value within the domesticated gene pool (Table S. 29) revealed that the Australian breeding selections (NMBPA) had the highest diversity and HAES cultivars had the lowest diversity (Figure S. 24). Other groups, such as HVP cultivars, heritage cultivars, cultivars bred by Norm Greber, cultivars bred by Backer and cultivars bred by Ian McConachie, show moderate to high diversity.

Moreover, results obtained for the cultivars and breeding selections grouped in clade I (n=4) in nuclear phylogenetic analysis performed in our previous publication showed an overall diversity of 1.13% across four genotypes. Individual diversity values ranging from 0.46% to 0.63% (Table S. 47) (Figure S. 25). The maximum number of accession-specific, unique, polymorphic SNP positions observed in Mac_279 (Table S. 48). Individual diversity for clade II genotypes (n=195) ranged from 0.4% to 1.13% (Table S. 49) (Figure S. 26). Cultivar ‘Young1’ (Mac_081) recorded the highest accession-specific, unique, polymorphic SNP positions (Table S. 50). A comparison of allelic diversity between two clades was present in Figure S. 27.

### Heterozygosity calculation

Among the wild *M. integrifolia* population, heterozygosity estimated based on the percentage of heterozygous SNP positions to reference genome size was the highest (0.71%) for Mac_052 and Mac_091 (Table S. 51). The majority of the values fell between 0.54% to 0.71%. A few outliers (Mac_250, Mac_171, Mac_058, Mac_163, Mac_139) with lower heterozygosity were also observed **(Fig. 1)**. For the wild *M. tetraphylla* population, highest heterozygosity value was recorded for Mac_236, and the lowest was for Mac_108 and MacP_15 (Table S. 52). Heterozygosity results showed that most of the values fall between 0.40% to 0.58% with a few outliers exhibiting both a lower (MacP_15, Mac_108, Mac_199, MacP_14, Mac_115, Mac_117, Mac_315) and a higher (Mac_238, Mac_236) heterozygosity level **(Fig. 2)**. *M. ternifolia* population, exhibited heterozygosity values between 0.35% to 0.42%, with only one genotype (Mac_126) showing a lower value, indicating the presence of a single outlier **(Fig. 3)**. The highest heterozygosity value of 0.42% was observed in Mac_300 (Table S. 53). *M. jansenii* population displayed heterozygosity values within the range from 0.27% to 0.34%, while three genotypes (Mac_216, Mac_217 and MacP_16) exhibited a reduced heterozygosity value thereby representing outliers in the distribution **(Fig. 4)** (Table S. 54). Domesticated macadamia genotypes showed most heterozygosity values between 0.33% and 0.95% except for a few outliers (Mac_350, Mac_348, Mac_333) at a higher level **(Fig. 5)** (Table S. 55).

**Fig. 1.**
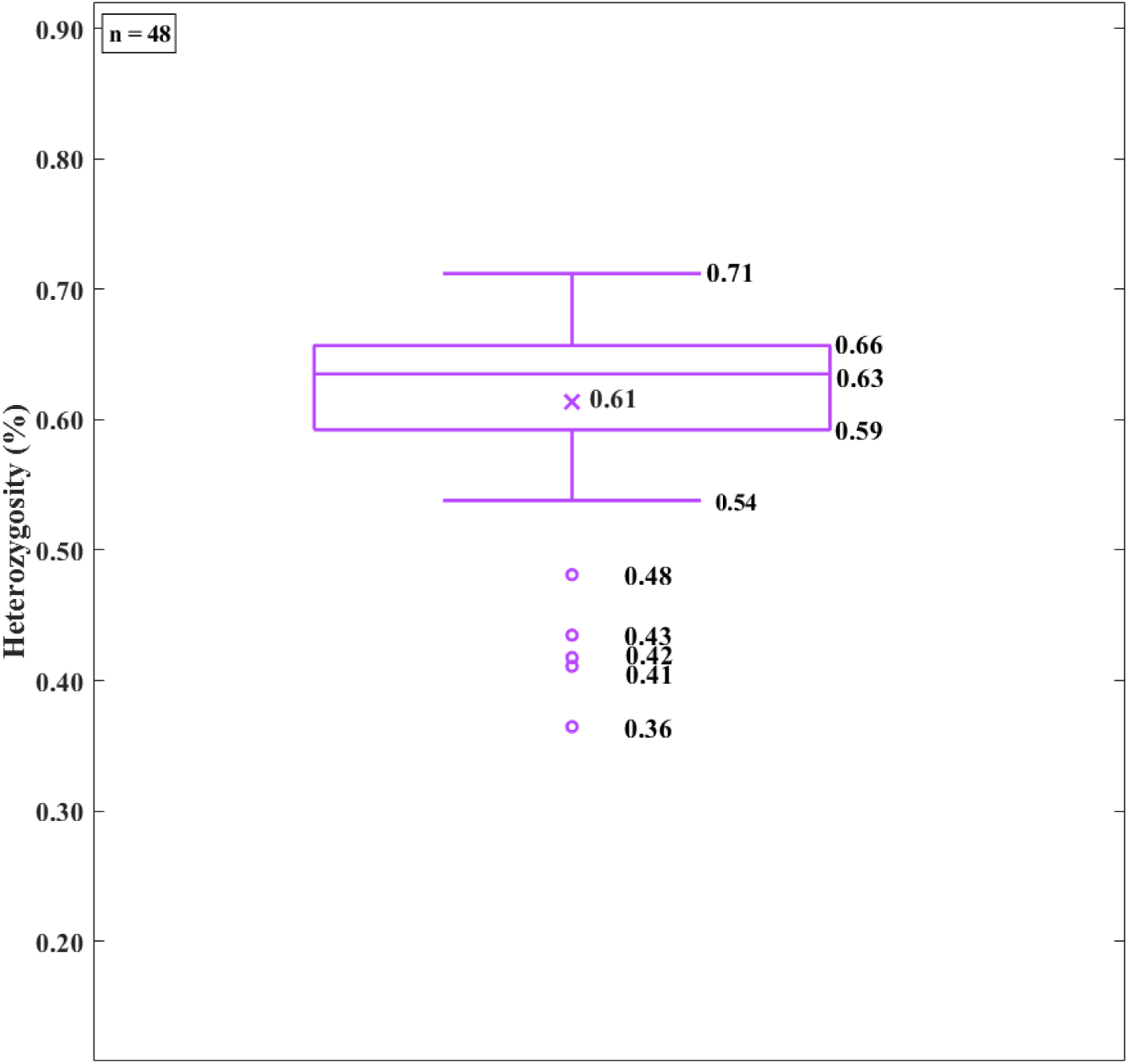
Distribution of heterozygosity across wild *M. integrifolia*.

**Fig. 2.**
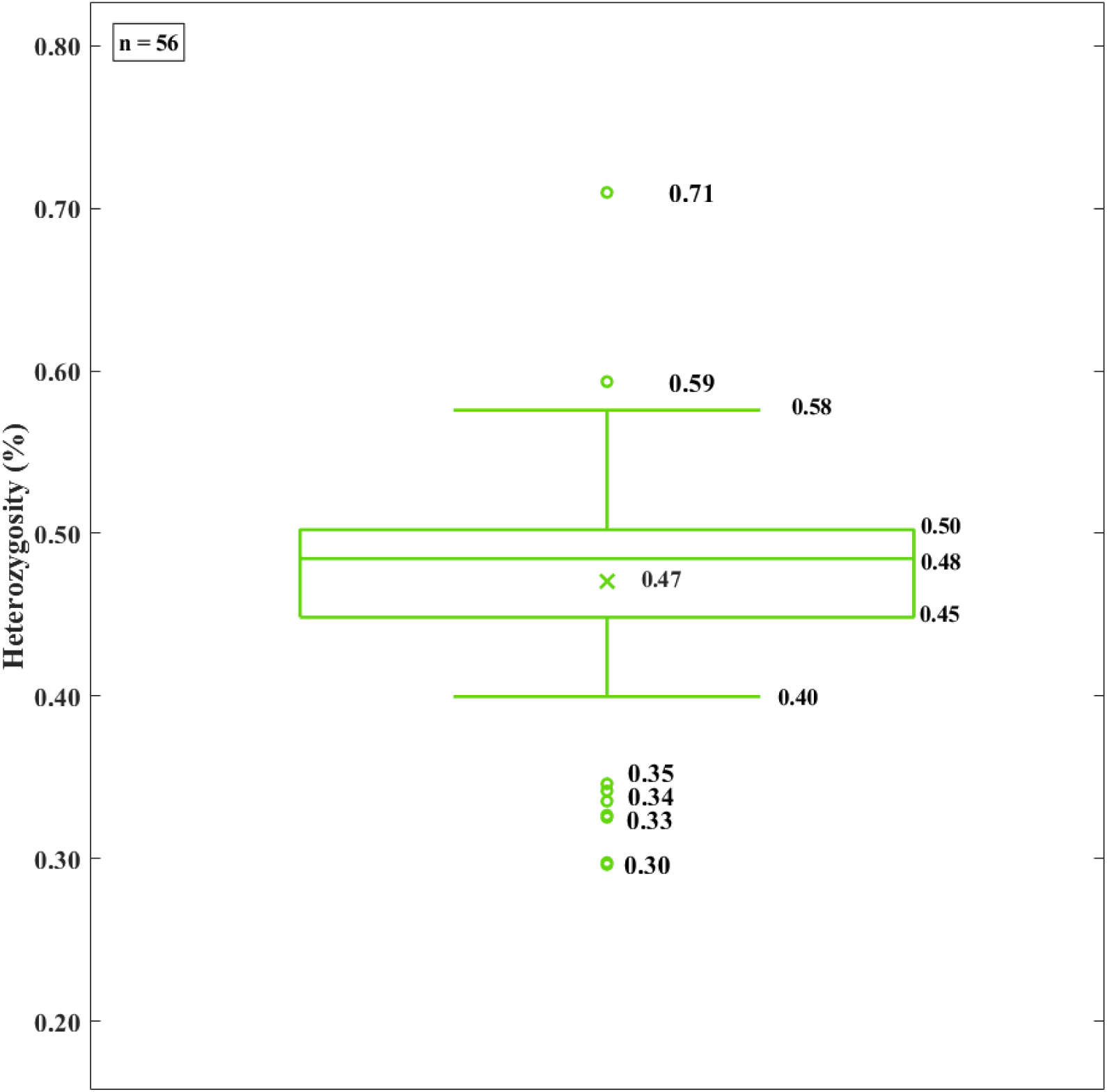
Distribution of heterozygosity across wild *M. tetraphylla*.

**Fig. 3.**
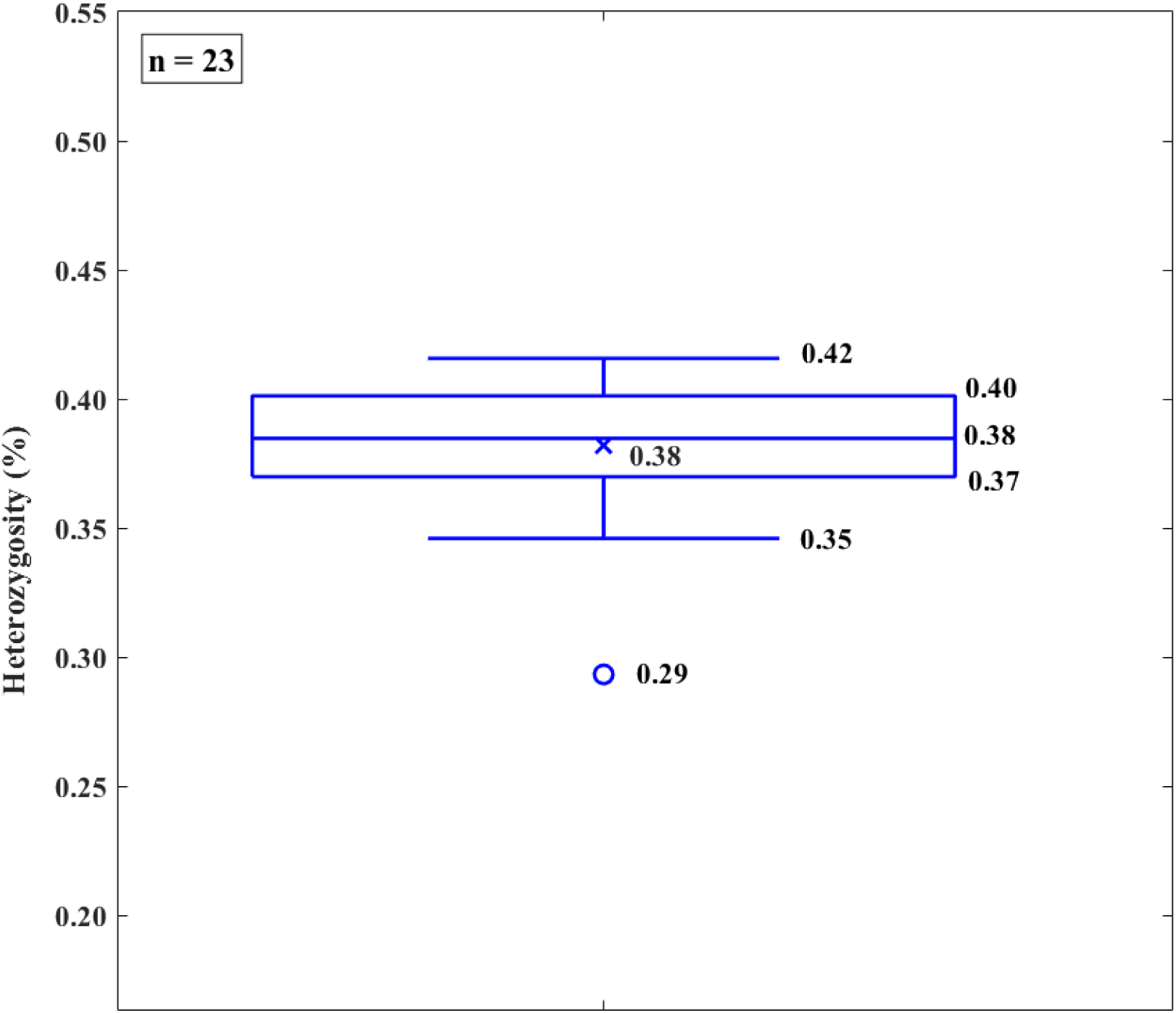
Distribution of heterozygosity across wild *M. ternifolia*.

**Fig. 4.**
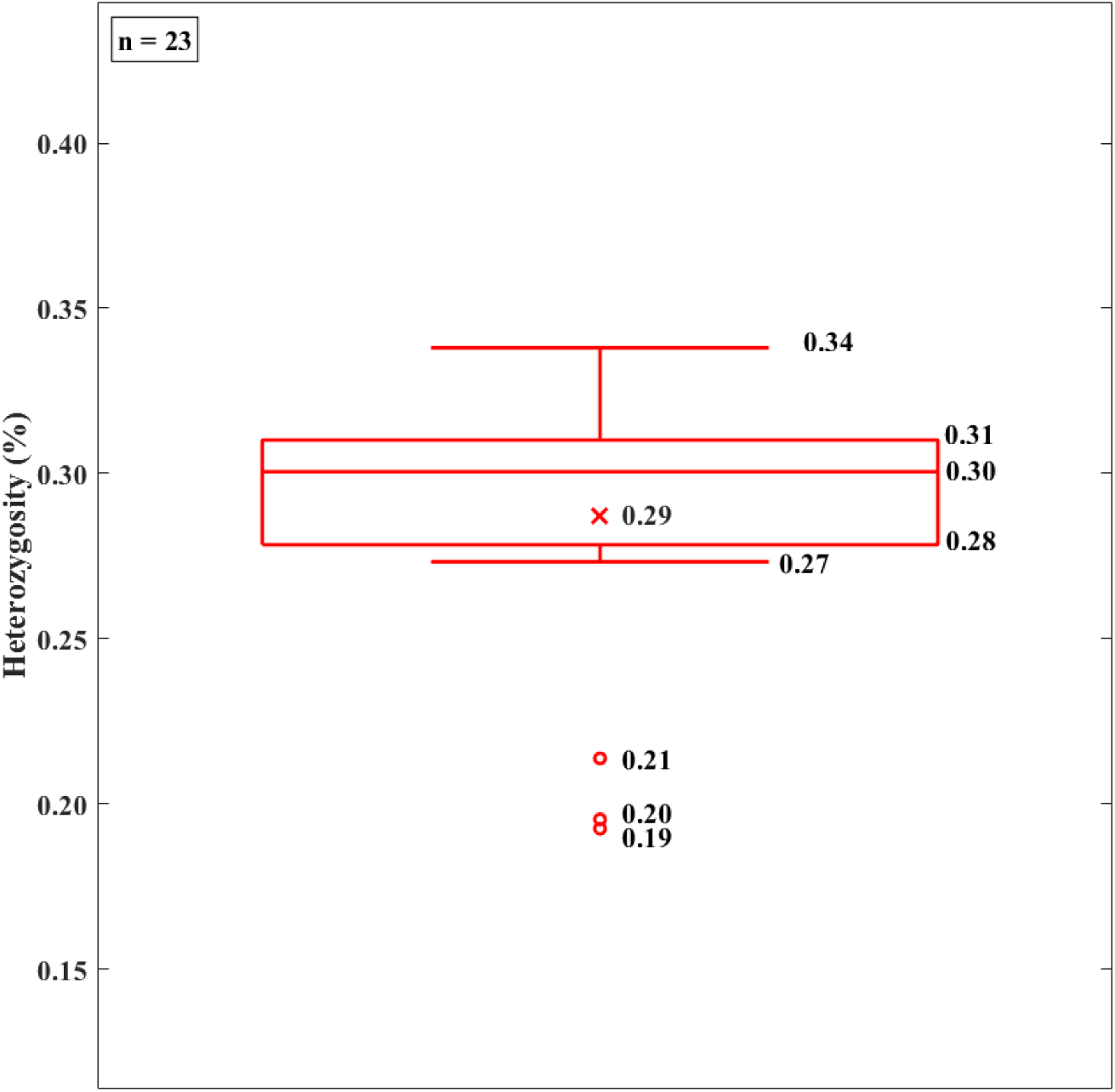
Distribution of heterozygosity across wild *M. jansenii*.

**Fig. 5.**
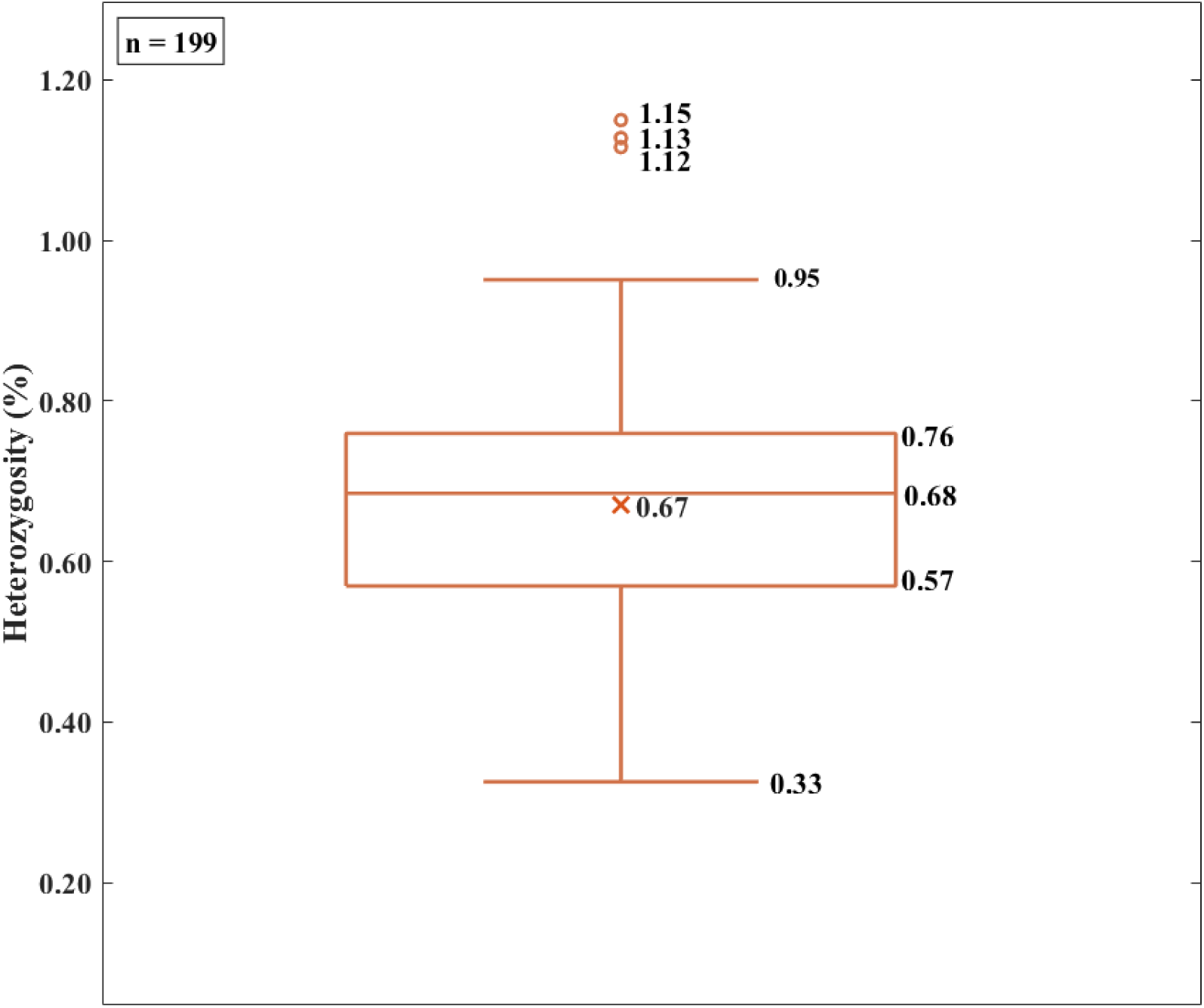
Distribution of heterozygosity across domesticated samples.

### Correlation between heterozygosity and phenotypic traits

Correlations were significant for three phenotypic traits (Figure S. 28). Among the assessed traits, yield efficiency (Kernel yield/tree volume) exhibited the strongest correlation with heterozygosity (r = 0.28, p < 0.05). Cumulative kernel yield at age 8 (r = 0.26, p < 0.05) and Cumulative NIS yield at age 8 (r = 0.25, p < 0.05) also showed a significant correlation with heterozygosity. However, no significant relationships were observed between heterozygosity and tree height at age 6, canopy volume at age 6 and percentage kernel recovery.

The relationship between the heterozygosity and yield efficiency (Kernel yield/tree volume) (Figure S. 29), heterozygosity and cumulative kernel yield at age 8 (Figure S. 30) and heterozygosity and cumulative NIS yield at age 8 (Figure S. 31) showed a nonlinear trend where yield efficiency or cumulative kernel yield at age 8 or cumulative NIS yield at age 8 increases with heterozygosity reaching an optimum level of performance and then declines beyond this point. The dotted curve line in Figure S. 29, S. 30 and S. 31 indicates this trend with a low coefficient of determination (R^2^) of 0.0887, 0.0897, and 0.0784, suggesting a weak but noticeable curvilinear relationship. Cluster 1 in Figures S. 29, S. 30 and S. 31 indicates genotypes with low heterozygosity and high performance, whereas cluster 2 indicates genotypes with high heterozygosity and low performance. In Figure S. 29, cluster 1, Mac_132, Mac_144, Mac_159, Mac_161, Mac_164, Mac_194 and Mac_198 exhibits low heterozygosity and high yield efficiency, whereas cluster 2 including Mac_082, Mac_186, Mac_230, Mac_280 and Mac_294, showed high heterozygosity and low yield efficiency. In Figure S. 30, the same set of genotypes clustered in cluster 1 of Figure S. 29 except for Mac_144 and Mac_161 exhibited low heterozygosity and high cumulative kernel yield, whereas cluster 2 consisting of Mac_010, Mac_048, Mac_088, Mac_096, Mac_176 along with the same genotypes presented in cluster 2 of Figure S. 29 exhibited high heterozygosity and low cumulative kernel yield. Low heterozygosity and high cumulative NIS yield were observed for five genotypes (Mac_132, Mac_159, Mac_164, Mac_194 and Mac_198) (Figure S. 31). High heterozygosity and low cumulative NIS yield was observed for Mac_082, Mac_186, Mac_230, Mac_280 and Mac_294. The non-significant traits (Height at age 6, canopy volume at age 6 and percentage of kernel recovery) showed a slight linear trend with heterozygosity (Figure S. 32).

## Discussion

The evaluation of genetic diversity has become a widely adopted approach for understanding wild germplasm and breeding materials, and helps conservation management and breeding programs [48, 49]. Many methods exist to evaluate diversity, based on percentage of polymorphic loci, polymorphism information content (PIC), mean number of alleles per locus and total gene diversity [6]. This study evaluated individual and population-level diversity based on the proportion of loci that display allelic variation within the population.

Allelic diversity was identified among domesticated genotypes from different breeding programs. The highest allelic diversity was exhibited in Australian breeding selections (NMBPA) compared to HVP cultivars, heritage cultivars, cultivars bred by Norm Greber, cultivars bred by Backer, cultivars bred by Ian McConachie, cultivars bred by Malawi and HAES cultivars. These findings align with the previous research based on SNPs identified through whole genome sequencing of accessions in a collection in China, which reported the highest diversity in Australian accessions (average number of nucleotide differences per site: Pi=1.818 x 10–3) and the lowest in Hawaiian accessions (Pi=1.496 x 10–3) [32]. Additionally, a study using SSR markers found that Australian Hidden Valley Plantation and pre-commercial selections had the second highest diversity (number of alleles per locus: Na = 8.46) following South Africa’s commercial and non-commercial local selections (Na = 9.31) [28].

This study also examined the allelic diversity at the population and individual genotype levels within the four wild populations. A previous study reported allelic diversity with markers at the population level for four wild species. It showed that *M. tetraphylla* exhibited the highest allelic diversity (Na = 15.73), followed by *M. integrifolia* (Na = 13.82), *M. ternifolia* (Na = 10.73) and *M. jansenii* (Na = 2.23) [29]. Another study also reported that *M. integrifolia* and *M. tetraphylla* exhibited higher diversity compared to *M. ternifolia* and *M. jansenii* [27]. The approach in this study enabled a more detailed understanding of the allelic diversity in genotypes collected from different natural distributions across Australia.

A previous study conducted for the northern, central and southern populations of *M. integrifolia* revealed a higher average number of nucleotide differences per site in the central population (n=15) (4.05 x 10−4), followed by the southernmost population (n=10) (2.87 x 10−4) and the least in the northern population (n=10) (2.76 x 10−4) based on SNP identified by GATK pipeline compared to the cultivar ‘Kau’ (HAES 344) reference genome [31]. This present study, found Mac_035 had the highest diversity based on the percentage of polymorphic loci. The observed difference could be due to the diversity calculation method, variant calling method, and the reference genome selection, which play a significant role in variant detection.

This study suggested that Mac_030, which is known as a planted tree, had a high diversity compared to the *M. tetraphylla* genome. The previous study identified Mac_030 as *M. tetraphylla* (corresponding to M259 in the previous study) [27]. However, our results showed Mac_030 had comparatively higher accession-specific unique polymorphic SNP positions (including both heterozygous and homozygous). Hence, additional analysis is needed to confirm its status as a pure *M. tetraphylla* accession.

Within the *M. ternifolia* population, we sequenced two representatives of the reference accession (Mac_309 and MacP_12). The results showed that MacP_12, the exact tree used for reference, showed the lowest diversity and had zero homozygous diverse positions, as expected.

In this study, the allelic diversity value reported for the individual *M. jansenii* genotypes was slightly different from the values reported previously [30]. We reported the individual diversity for the same eight genotypes. Among the eight genotypes, six (MacP_01, MacP_02, MacP_4, MacP_05, MacP_06 and MacP_07) exhibited slightly higher diversity than that reported in the previous study. The remaining two genotypes (MacP_03 and MacP_16) reported lower diversity values. Overall diversity in the present study (1.10%) was higher than the value reported previously (0.92%), which may be due to the size of the population, which significantly influences diversity analysis [27, 50]. Moreover, the present diversity pipeline was structured to remove the data noise, which can lead to the identification of erroneous diverse positions. Erroneous diverse positions were identified and removed from the total diverse positions [45]. This could be another reason for the observed difference between the current analysis and the previously reported values for the *M. jansenii* population [30].

The results of this study showed for the first time a difference in allelic diversity between wild *M. integrifolia* and domesticated *M. integrifolia*. These findings highlight the need to protect wild material because it holds essential genetic diversity, which will serve as a foundation for future breeding programs.

Moreover, 19 genotypes identified with zero number of accession-specific, unique, polymorphic homozygous SNP positions compared to the reference genome could be due to the closely relatedness, but need further validation in future. Furthermore, in future, the alleles identified in this study can be used to screen for novel alleles in wild macadamia that are not present in the domesticated material, which can be utilized in future breeding programs.

Heterozygosity in plants has been a subject of extensive research and is commonly used to compare genetic variation within populations and between different populations. It is commonly reported as observed and expected heterozygosity, where expected heterozygosity relies on allele frequency and observed heterozygosity comes directly from individual genotypic information and has been reported for many plants using different molecular markers [33]. Accurate measurement of heterozygosity is very crucial to understand the structure of a population. This study reports the genome-wide or autosomal heterozygosity, which is known to be more comprehensive and unbiased by sample size. Genome-wide heterozygosity estimates a proportion of heterozygous sites across all the nucleotide positions, which consist of both polymorphic and monomorphic sites [33].

In the present study, heterozygosity in domesticated material ranged from 0.33% to 1.15%, while heterozygosity in wild *M. integrifolia*, *M. tetraphylla*, *M. ternifolia* and *M. jansenii* ranged from 0.36% to 0.71%, 0.30% to 0.71%, 0.29% to 0.42% and 0.19% to 0.34% respectively. The heterozygosity values observed in the current study fall within a similar range to those reported for walnut 0.385% [51], hazelnut 0.334% to 0.855% [52], and cassava (0.61 – 0.84% [53]. In contrast, crops like pistachio (1.2%) [54] and mango 2.5% [55] display substantially higher heterozygosity values. For macadamia, a previous study reported on the heterozygosity of all four species [29]. It was reported that the highest heterozygosity was in *M. tetraphylla* (He = 0.76), followed by *M. integrifolia* (He = 0.73), *M. ternifolia* (He = 0.72) and he lowest was in *M. jansenii* (He = 0.47). In contrast, the current study showed that the range of heterozygosity of the two edible species is comparable. Additionally, the present results for *M. jansenii* were highly consistent with the previous findings that reported a low level of heterozygosity based on genome-wide heterozygosity [30].

This analysis revealed a wide range of heterozygosity in the domesticated macadamia and showed that several cultivars and breeding selections had higher levels of heterozygosity than wild materials, even though it is generally reported that wild plant populations exhibit greater genetic heterozygosity [56]. Domestication usually reduces heterozygosity due to loss of diversity via selection. However, with macadamia domestication there has been interspecific hybridization which has increased heterozygosity. Previously a number of studies have reported the heterozygosity for cultivars and breeding selections [23, 26, 28, 29, 32, 36–38, 41]. The present study showed both accessions of *M. integrifolia* 741 exhibited low heterozygosity values (0.40 and 0.41) compared to the previously reported value (0.98%) [38]. Furthermore, the current study showed highest heterozygosity for the Australian breeding selections (Mac_333, Mac_348, Mac_350) followed by cultivar A268 (Mac_280) which is known as a hybrid of *M. integrifolia* and *M. tetraphylla* [41]. However, the heterozygosity value reported for A268 is higher than the previously reported value of 0.18 based on SNP markers [41]. The previously reported value of 0.16 for cultivar A4 was also found to be higher in this study. According to Hardenbol *et al.,* a possible explanation could be when using codominant markers such as SNPs and SSRs, only one of the two allele is detected hence heterozygosity is under calculated due to null alleles [41, 57]

The relationship between heterozygosity at the individual level and the phenotypic traits has been examined in both plant and animal species [58–61]. Previous studies have reported that there was no relationship between heterozygosity and kernel recovery and yield efficiency, but have shown a significant relationship with cumulative kernel yield, canopy volume, cumulative nut in shell yield yield and tree height at age 6 in macadamia [26]. This study supports the previous findings of kernel recovery where no significant relationship is observed with heterozygosity [26]. However, the current findings show a significant relationship with yield efficiency but no significant relationship with tree height or canopy volume. Both studies reported a significant relationship between heterozygosity and the cumulative NIS yield. The result of the current study indicates there is an optimum level of heterozygosity related to yield performance. Heterozygosity above this optimum level may result in reduced performance. Hence, selecting highly heterozygous cultivars or breeding selections is not necessarily desirable; selecting those with an optimum level would be more appropriate. To ensure that our results were not due to lack of sequence reads, potential causing a failure to capture heterozygous SNPs and thus reducing heterozygosity in the genotypes in cluster 1, we compared the mapping percentage of the genotypes in cluster 1 and 2 and found no difference between them (Figure S. 33).

## Conclusions

This study presents the first report of genome-wide heterozygosity at the individual level for both wild and domesticated macadamia genotypes. The availability of SNP data and accession-specific unique SNPs compared to the reference for individual genotypes for wild and domesticated material facilitates marker development in future breeding programs. Assessment of the relationship between heterozygosity and phenotypic traits revealed that selecting cultivars or breeding selections with an optimum level of heterozygosity would be more appropriate than selecting cultivars with higher heterozygosity. This information proved to be a valuable genomic resource for use to improve future breeding programs.

## Acknowledgements

We thank Mr Rod Daley for collecting Phenotypic data and A/Prof Craig Hardner for analyzing phenotypic data. We also thank The University of Queensland Research Computing Centre (UQ-RCC) for providing all the computation resources, and Denise Bond from Macadamia Conservation Trust for sharing data.

## Contribution

**RH:** Design, Methodology, Supervision, Project administration, Funding acquisition, Data Curation, Resources and Writing - Review & Editing. **AF**: Design, Methodology, Supervision, Project administration, Formal analysis, Resources, Data Curation, Writing - Review & Editing. **BT**: Design, Methodology, Supervision, Project administration, Data Curation, Writing - Review & Editing. **MA**: Methodology, Supervision, Data Curation, Writing - Review & Editing. **PM**: Methodology, Writing - Review & Editing. **SM**: Methodology, Formal analysis, Writing an original draft.

## Data availability

All sequence data for wild material are available at NCBI under BioProject PRJNA1036028 (BioSample numbers SAMN38356272–SAMN38356438), and for domesticated material under BioProject PRJNA1089341 (BioSample numbers SAMN40990754–SAMN40990952).

## List of abbreviations

HAES: Hawaii Agricultural Experiment Station
HVP: Hidden Valley Plantations
Indels: Insertions and deletions
MNPs: Multi-nucleotide polymorphisms
NIS: Nut in shell
NMBPA: Australian macadamia breeding program accession
NSW: New South Wales
PIC: Polymorphism information content
SNPs: Single-nucleotide polymorphisms
SSR: Simple sequence repeat

---

**Figure S. 1.**
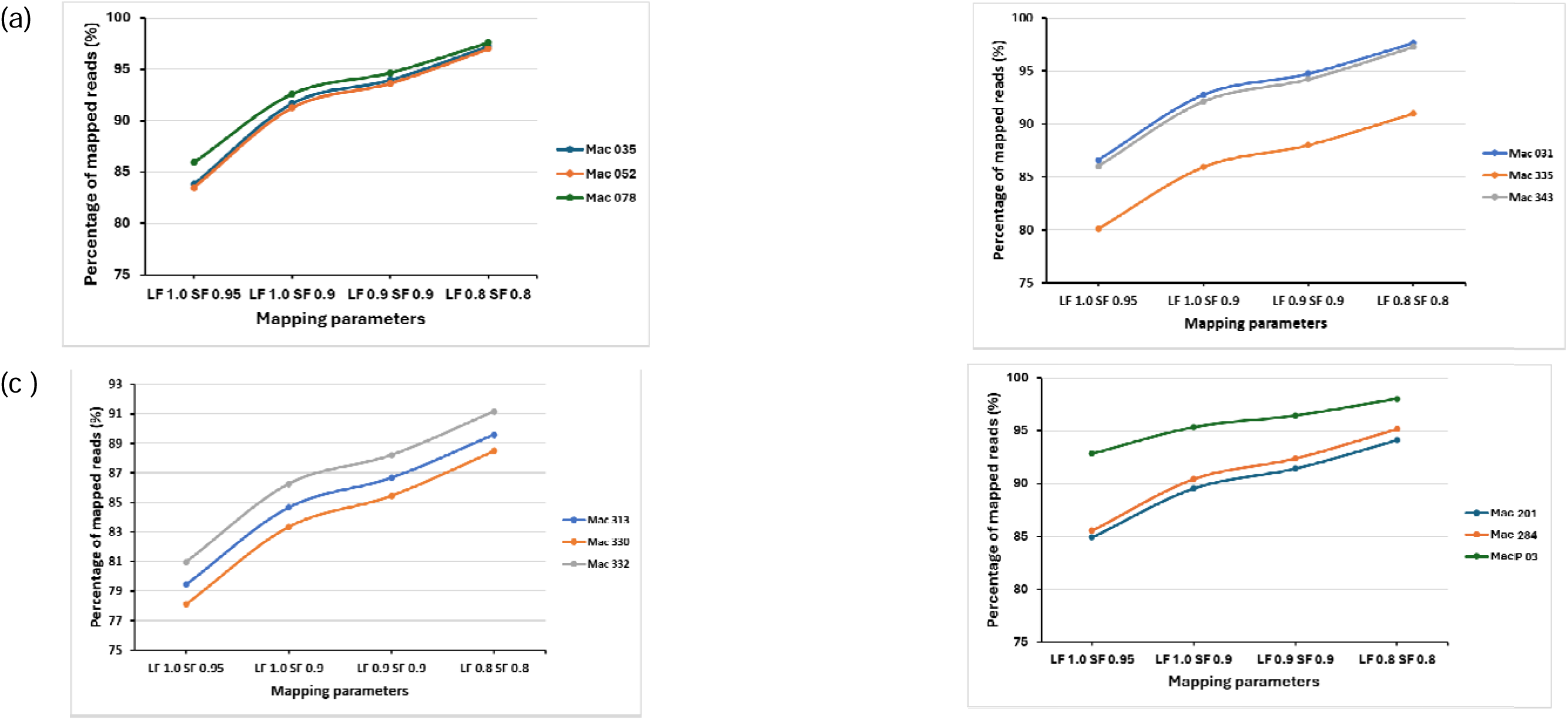
The mapping titration performed at four different stringency levels of read length fraction (LF) and read similarity fractions (SF) (LF 1.0 SF 0.95, LF 1.0 SF 0.9, LF 0.9 SF 0.9, LF 0.8 SF 0.8) (a) Mapping titration of three *M. integrifolia* genotypes against *M. integrifolia* reference genome (b) Mapping titration of three *M. tetraphylla* genotypes against *M. tetraphylla* reference genome. (c) Mapping titration of three *M. ternifolia* genotypes against *M. ternifolia* reference genome. (d) Mapping titration of three *M. jansenii* genotypes against *M. jansenii* reference genome

**Figure S. 2.**
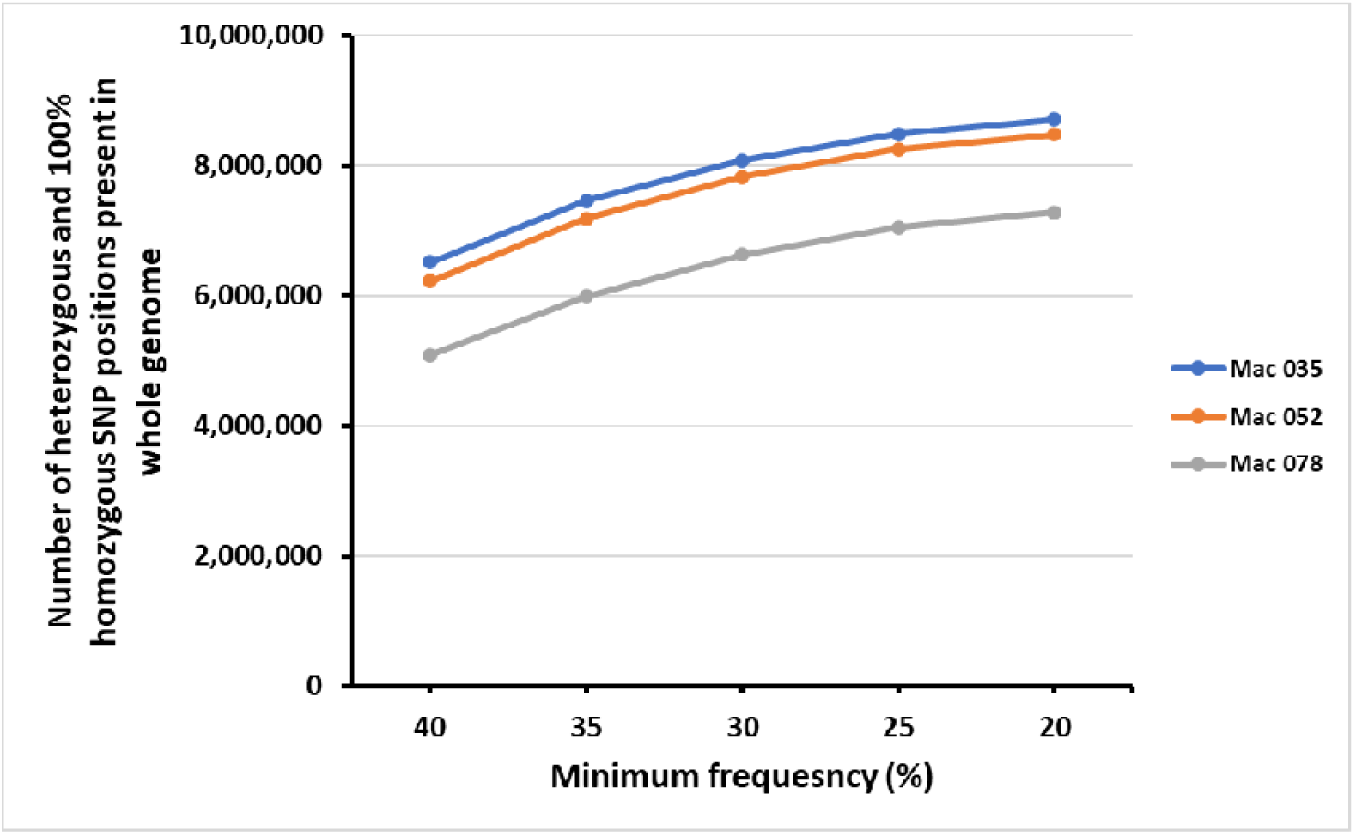
Frequency titration. Frequency titration was performed at five levels of minimum frequency percentage (40%, 35%, 30%, 25% and 20%).

**Figure S. 3.**
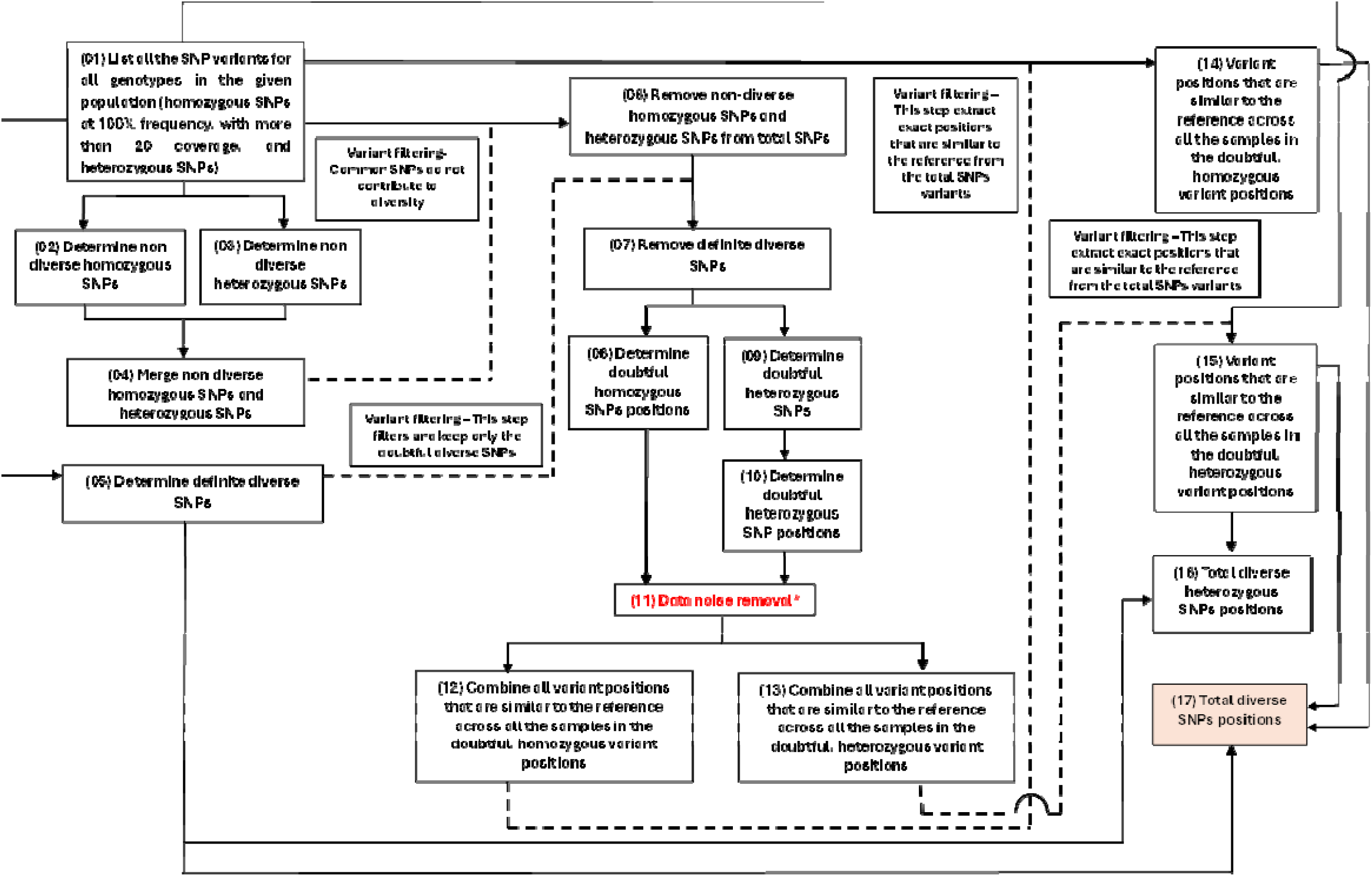
Diversity analysis pipeline

**Figure S. 4.**
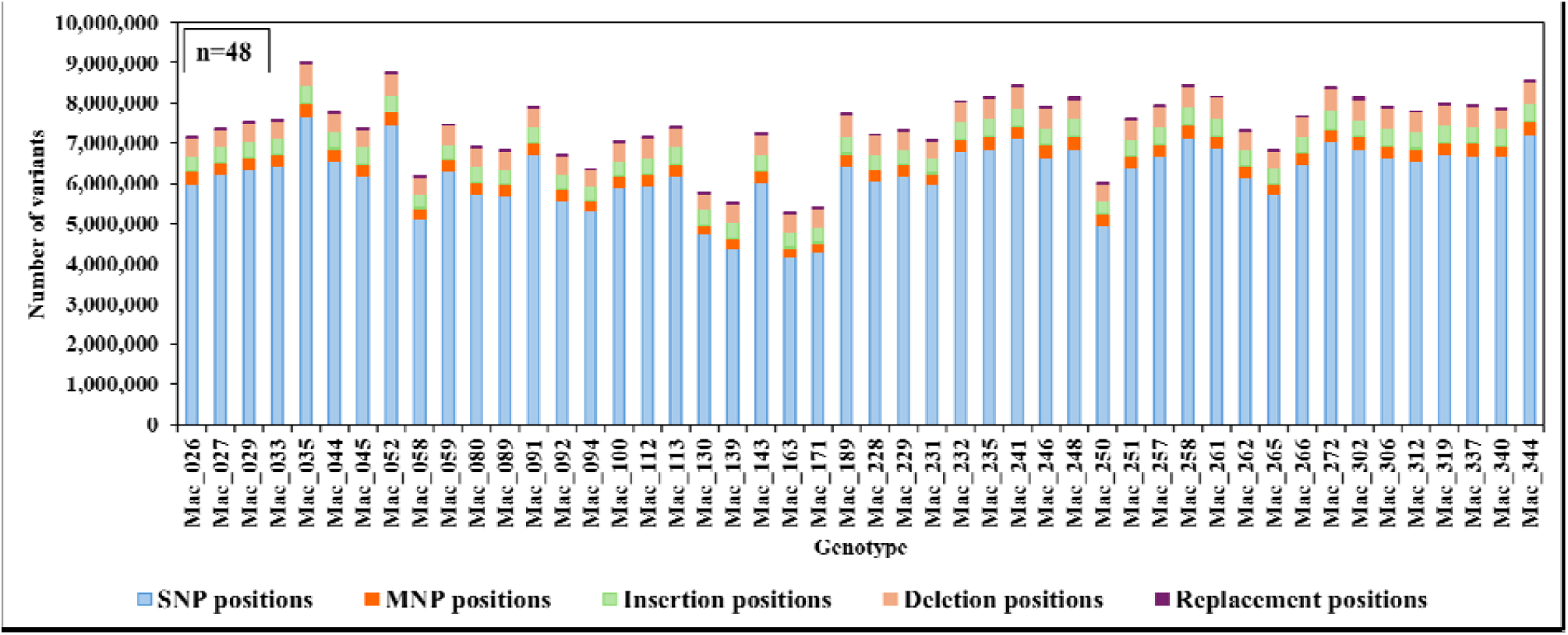
Number of variants in the whole genome across wild *M. integrifolia*.

**Figure S. 5.**
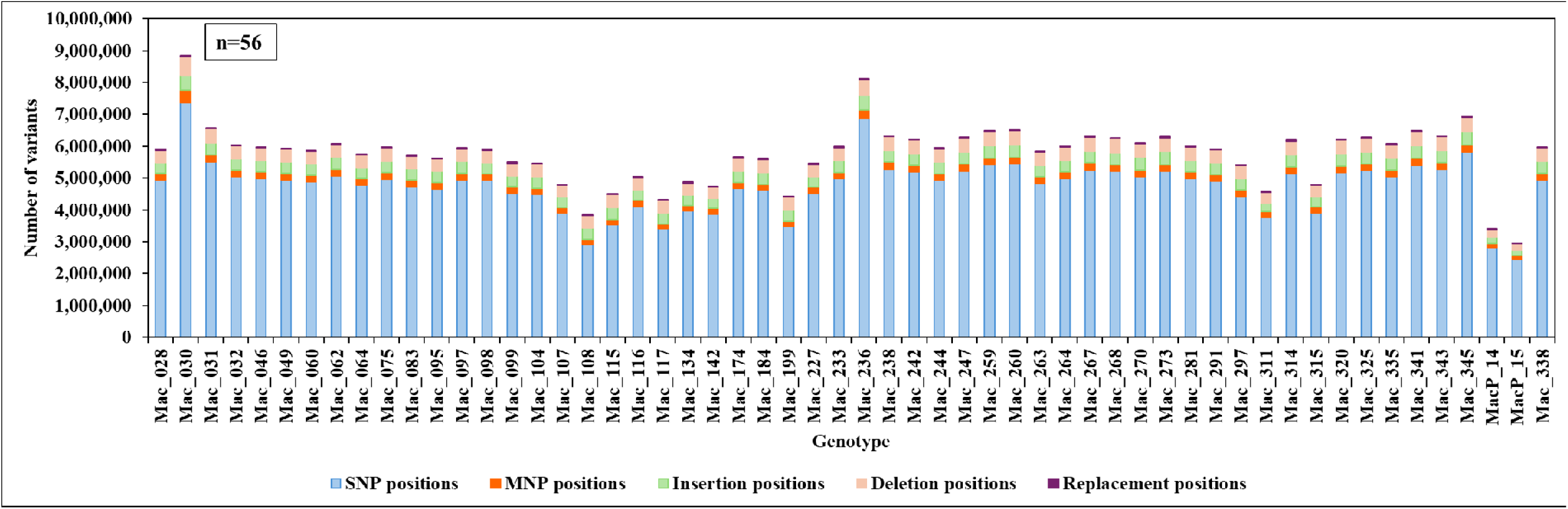
Number of variants in the whole genome across wild *M. tetraphylla*.

**Figure S. 6.**
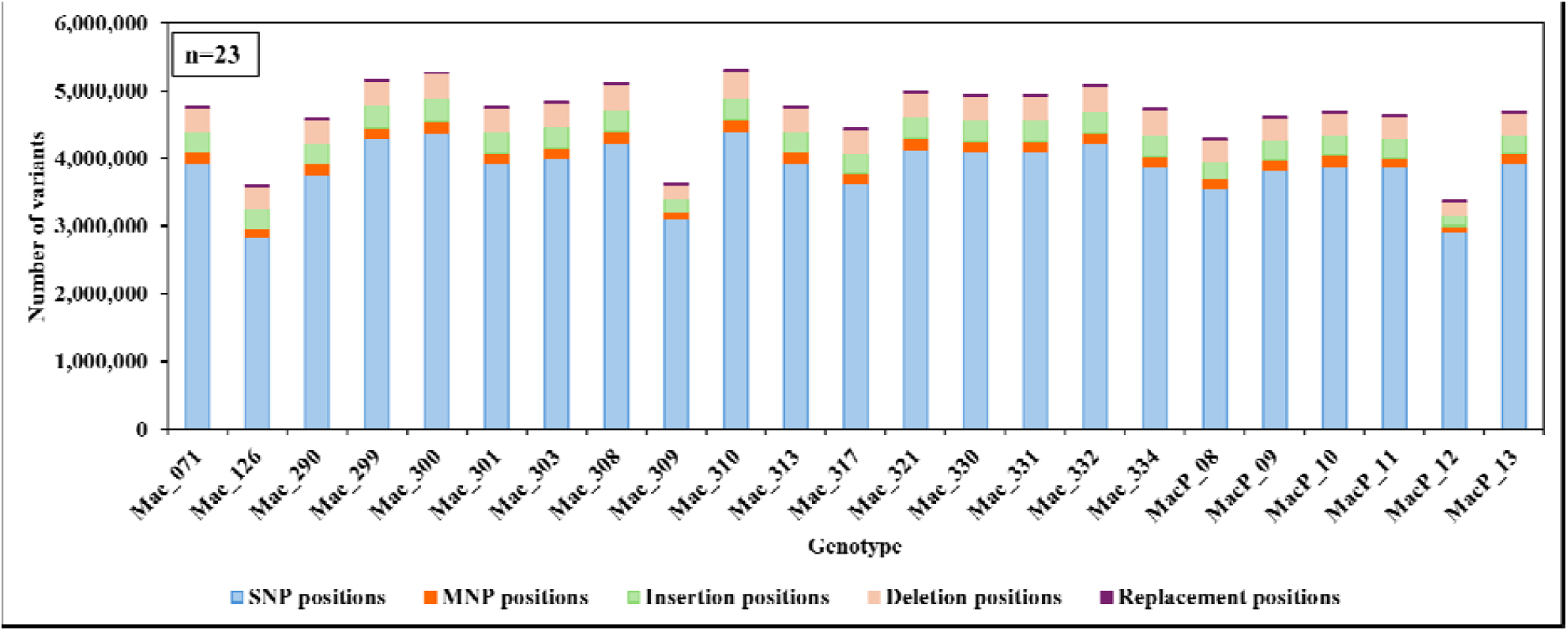
Number of variants in the whole genome across wild *M. ternifolia*.

**Figure S. 7.**
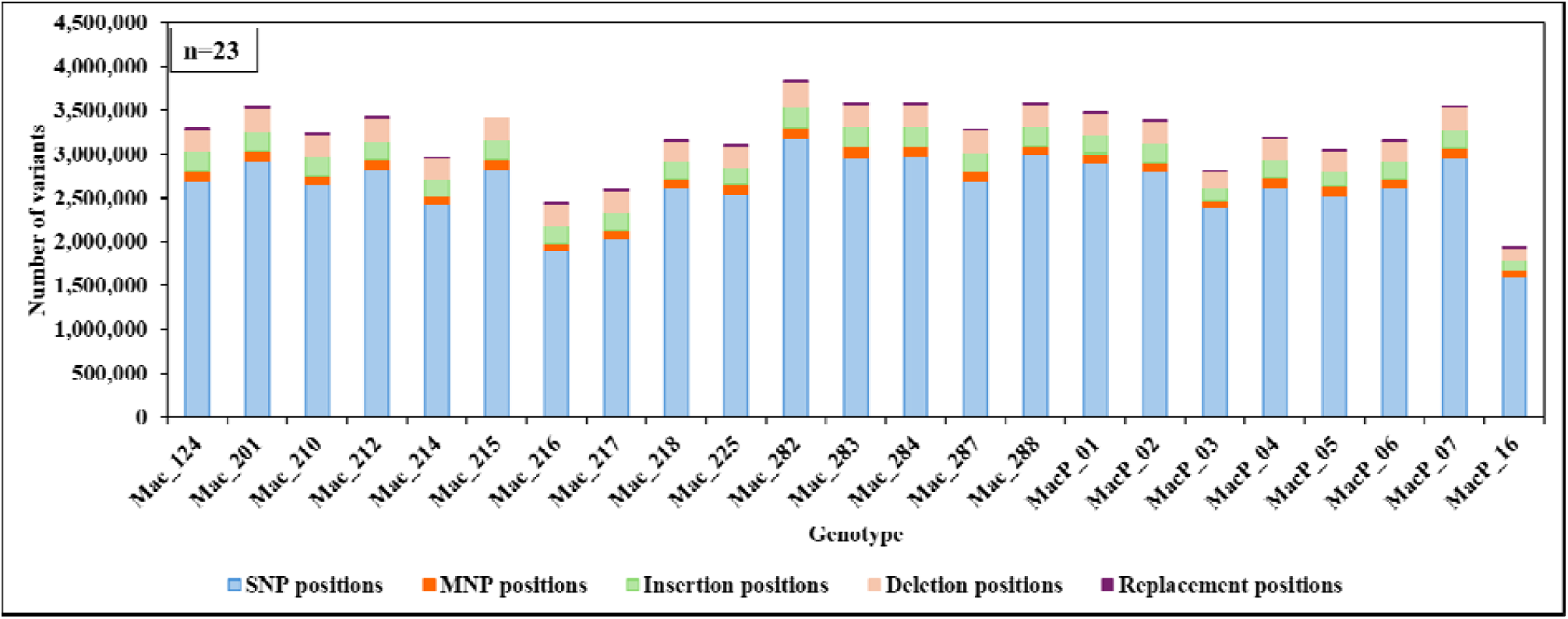
Number of variants in the whole genome across wild *M. jansenii*.

**Figure S. 8.**
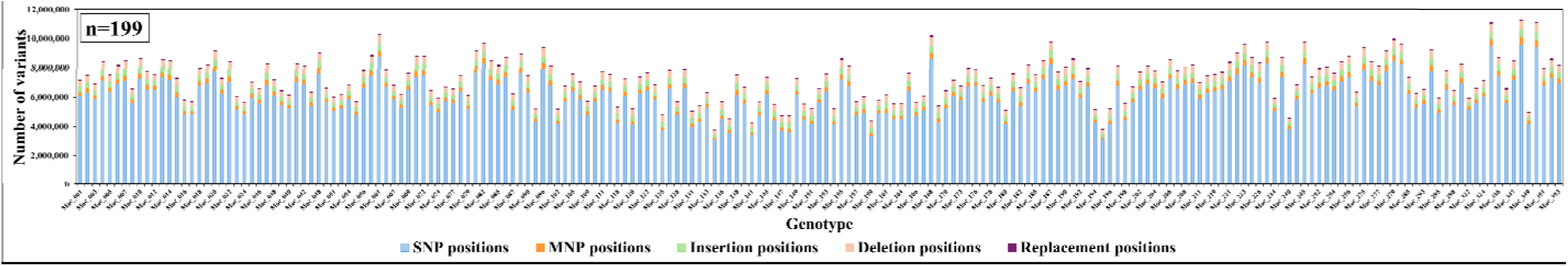
Number of variants in the whole genome across domesticated samples.

**Figure S. 9.**
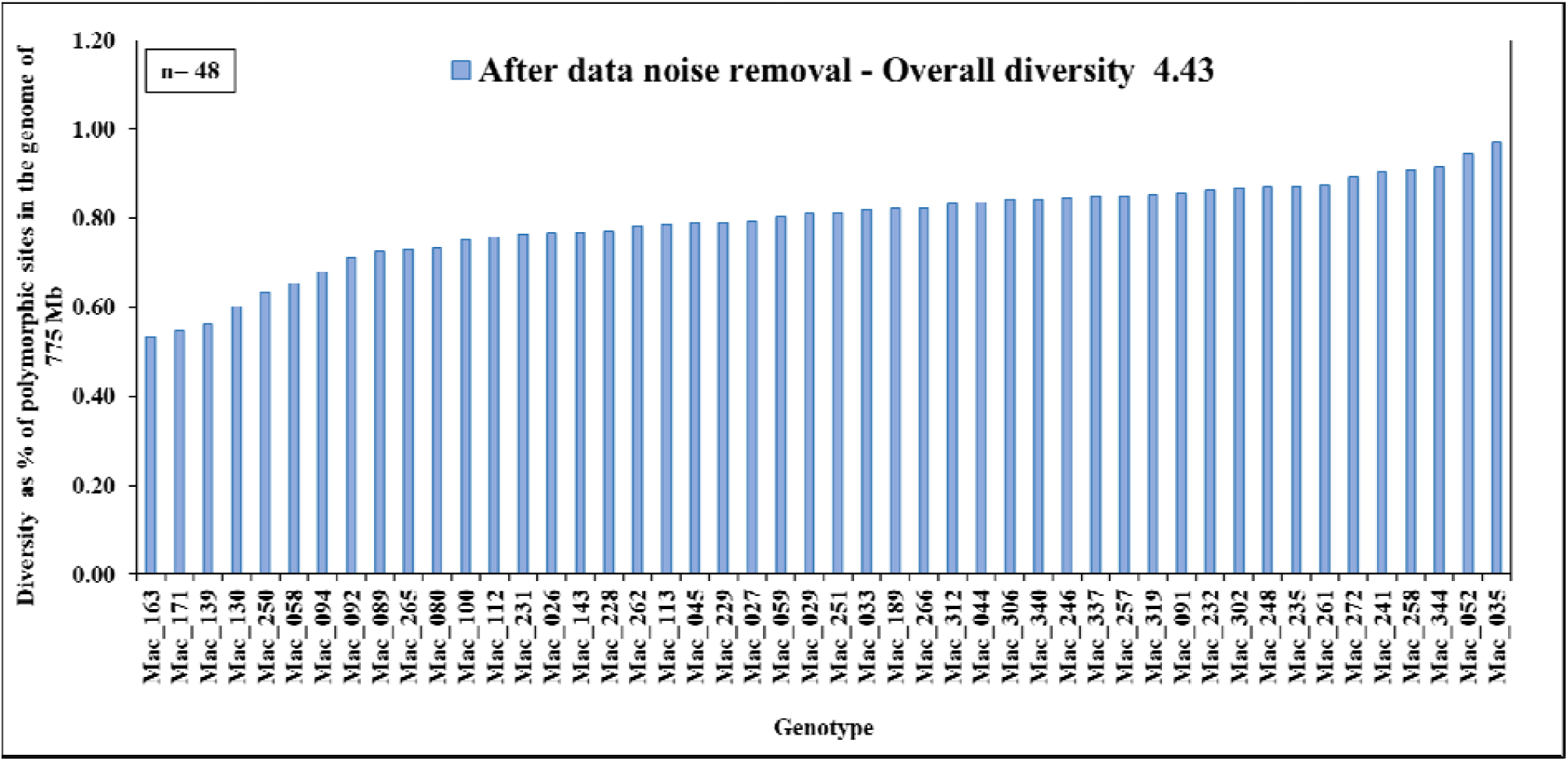
Diversity within wild *M. integrifolia*. Diversity was calculated as a percentage of polymorphic sites in the genome of *M. integrifolia* (775 Mb)

**Figure S. 10.**
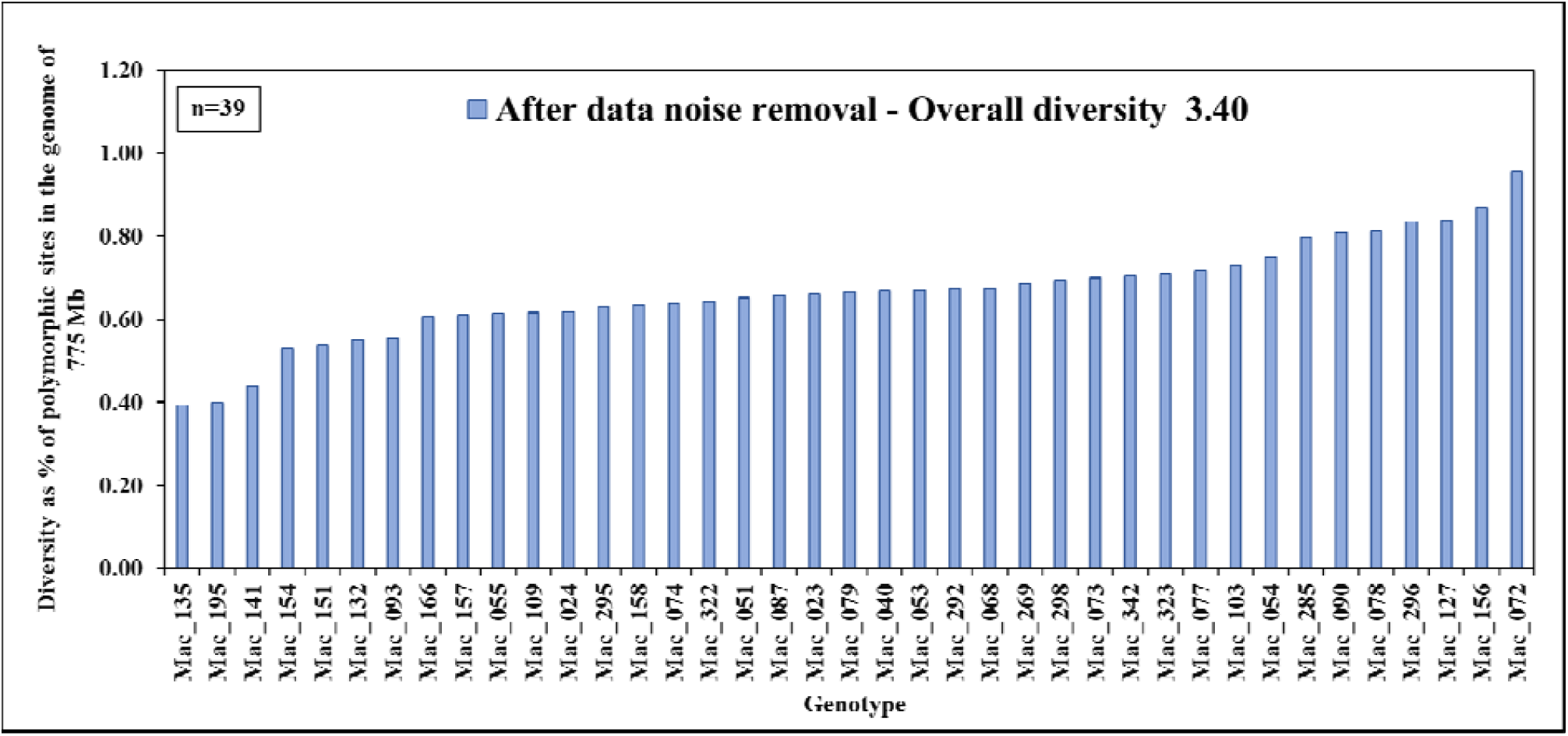
Diversity within domesticated *M. integrifolia*. Diversity was calculated as a percentage of polymorphic sites in the genome of *M. integrifolia* (775 Mb)

**Figure S. 11.**
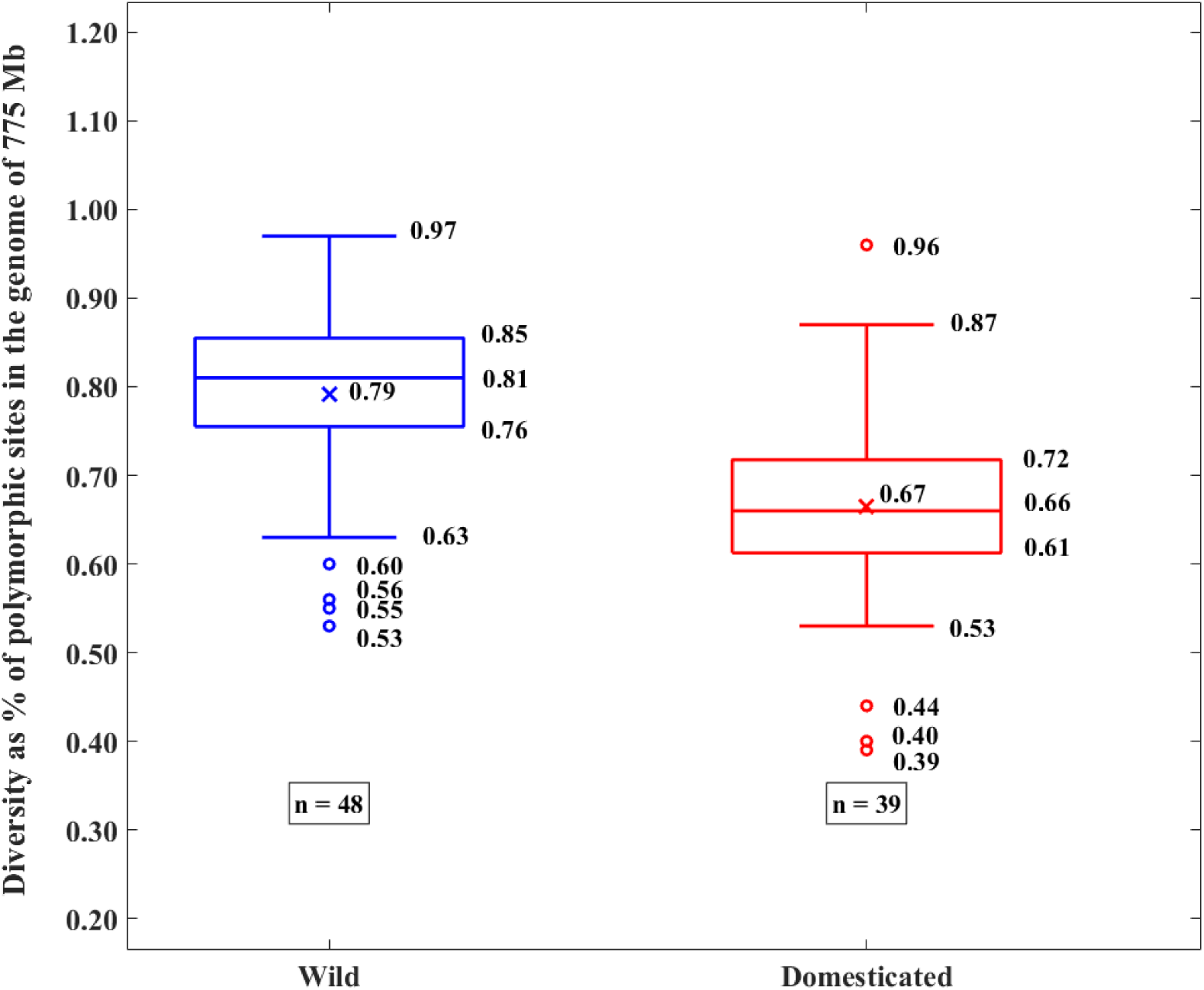
Comparison of diversity between wild and domesticated *M. integrifolia*. Diversity was calculated as a percentage of polymorphic sites in the genome of *M. integrifolia* (775 Mb)

**Figure S. 12.**
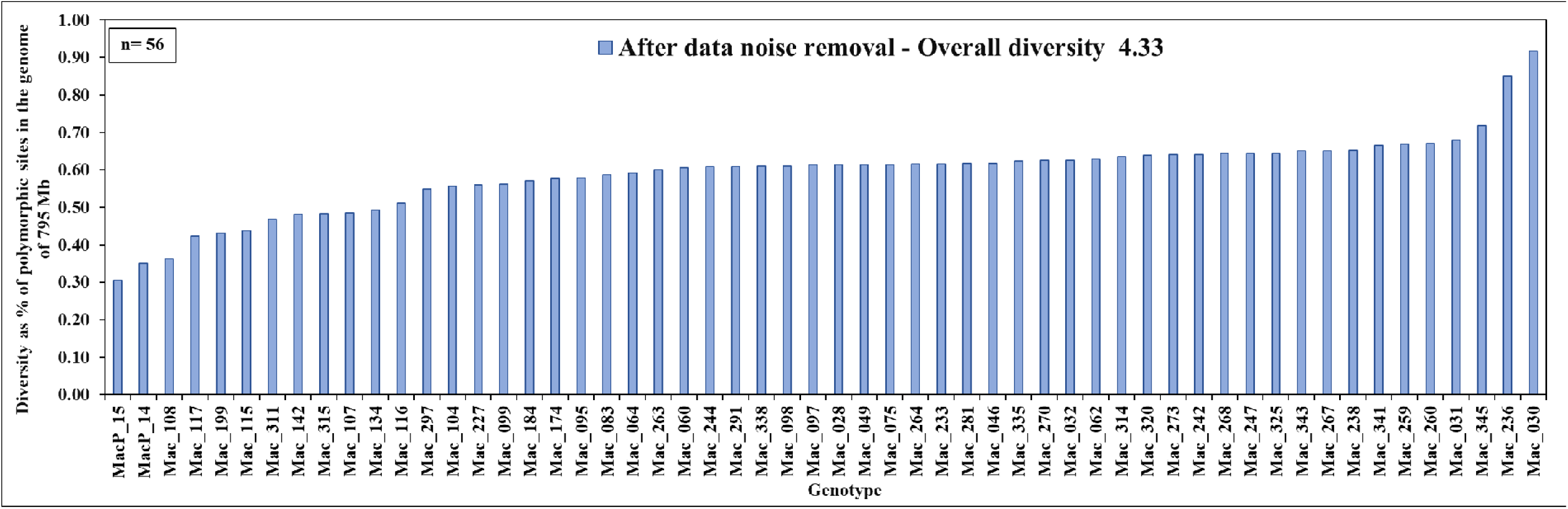
Diversity within wild *M. tetraphylla*. Diversity was calculated as a percentage of polymorphic sites in the genome of *M. tetraphylla* (795 Mb)

**Figure S. 13.**
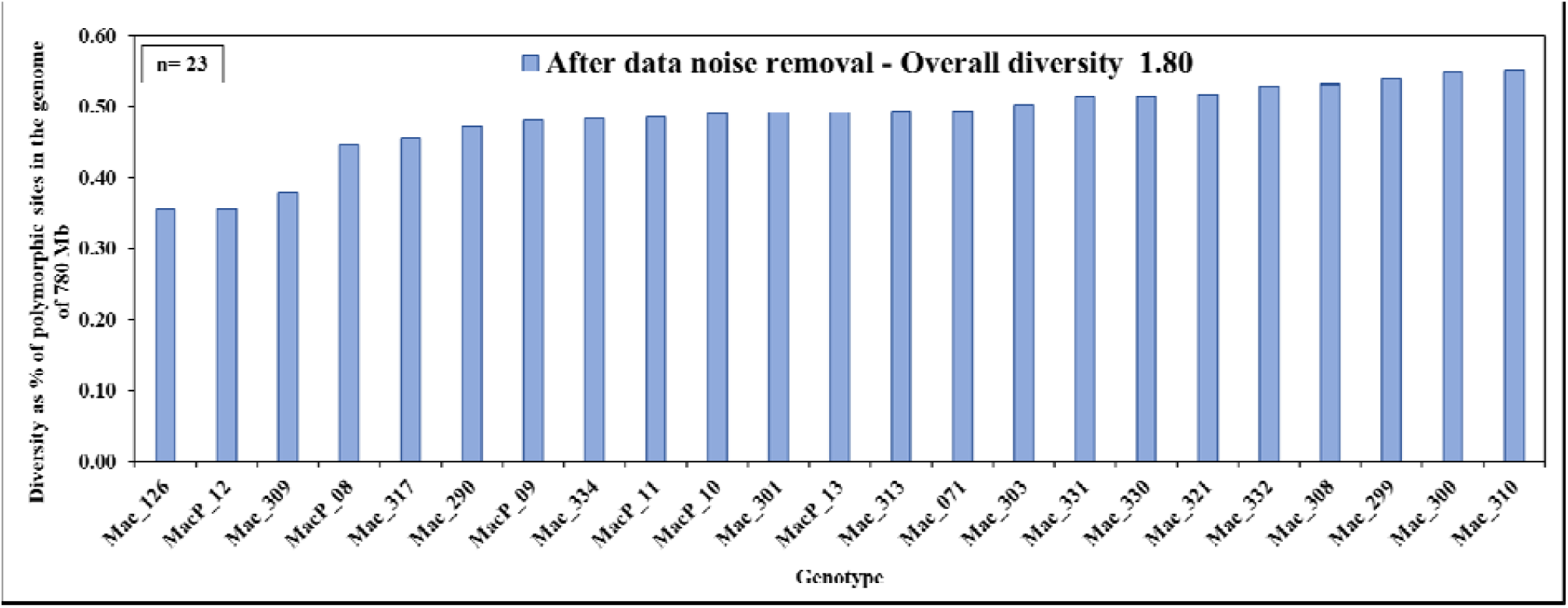
Diversity within wild *M. ternifolia*. Diversity was calculated as a percentage of polymorphic sites in the genome of *M. ternifolia* (780 Mb)

**Figure S. 14.**
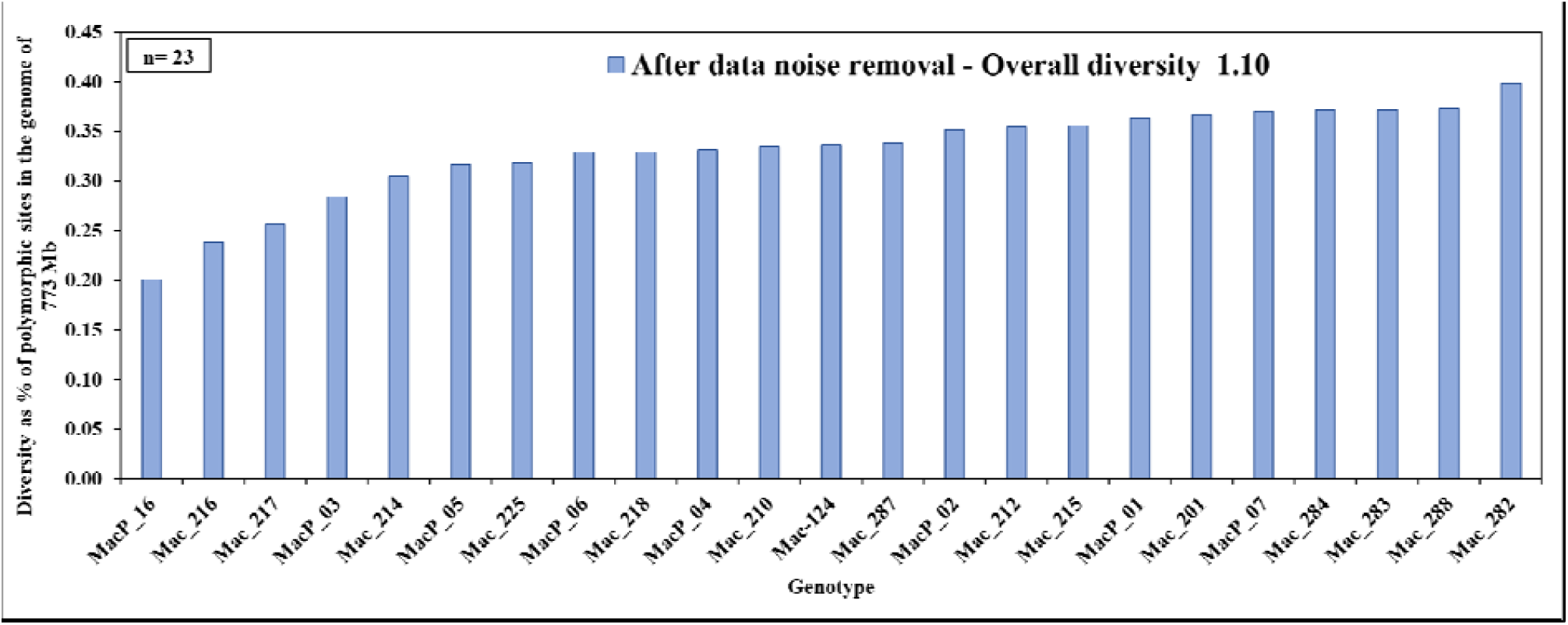
Diversity within wild *M. jansenii*. Diversity was calculated as a percentage of polymorphic sites in the genome of *M. jansenii* (773 Mb)

**Figure S. 15.**
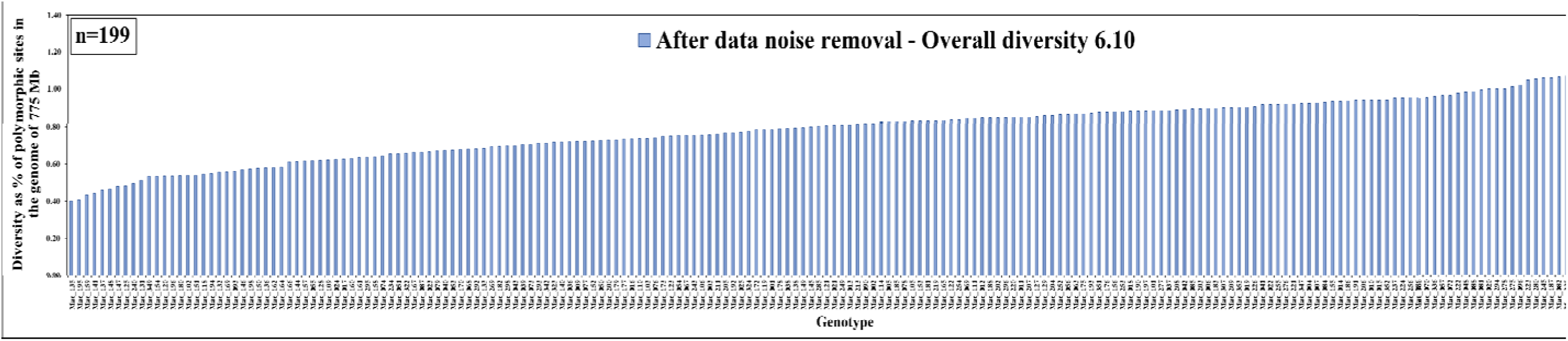
Diversity within domesticated cultivars and breeding selections. Diversity was calculated as a percentage of polymorphic sites in the genome of *M. integrifolia* (775 Mb)

**Figure S. 16.**
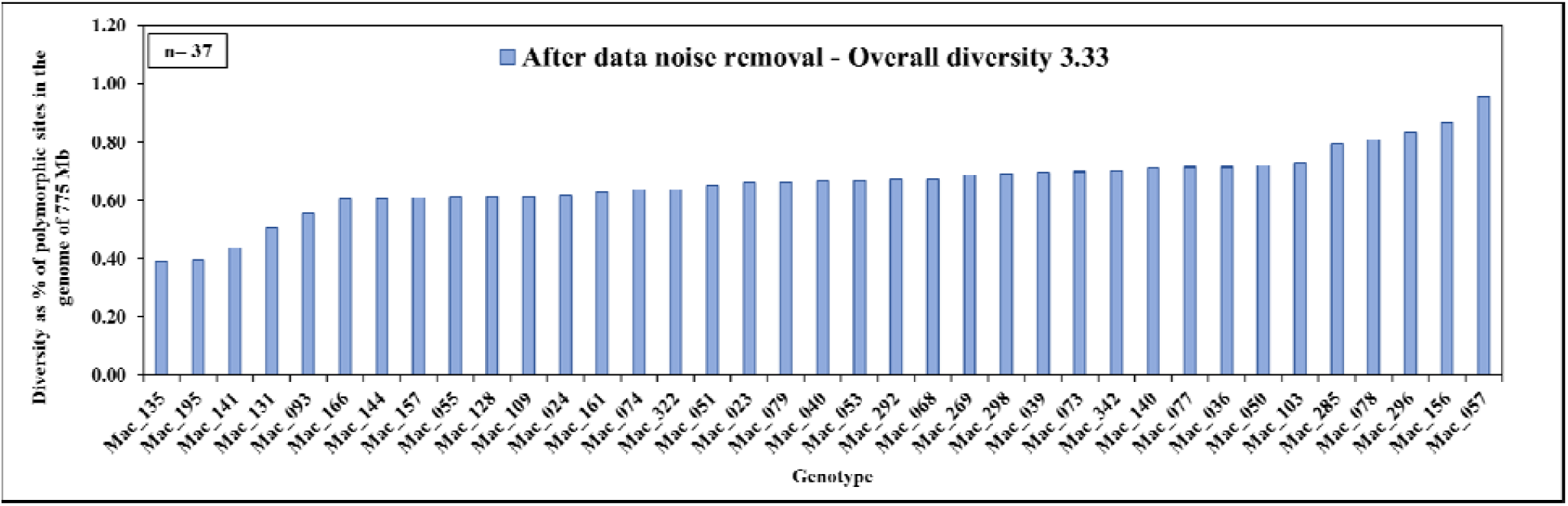
Diversity within Hawaiian Agricultural Experiment Station (HAES) cultivars. Diversity was calculated as a percentage of polymorphic sites in the genome of *M integrifolia* (775 Mb)

**Figure S. 17.**
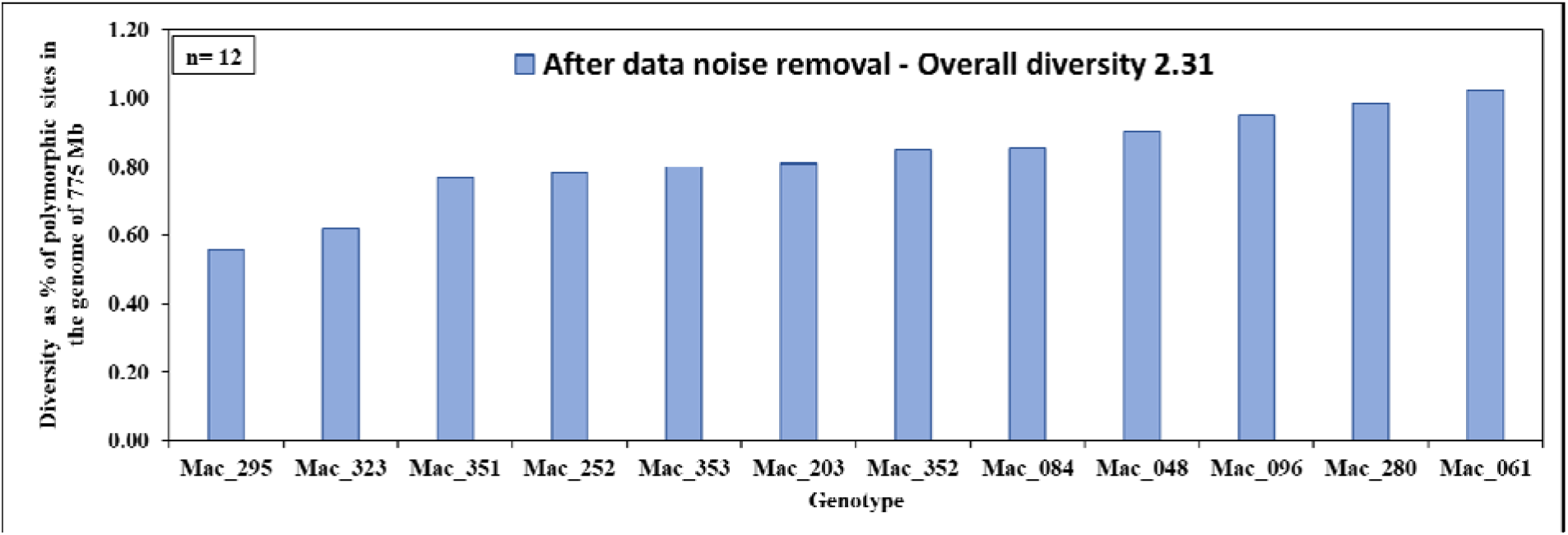
Diversity within Hidden Valley Plantation cultivars (HVP). Diversity was calculated as a percentage of polymorphic sites in the genome of *M. integrifolia* (775 Mb)

**Figure S. 18.**
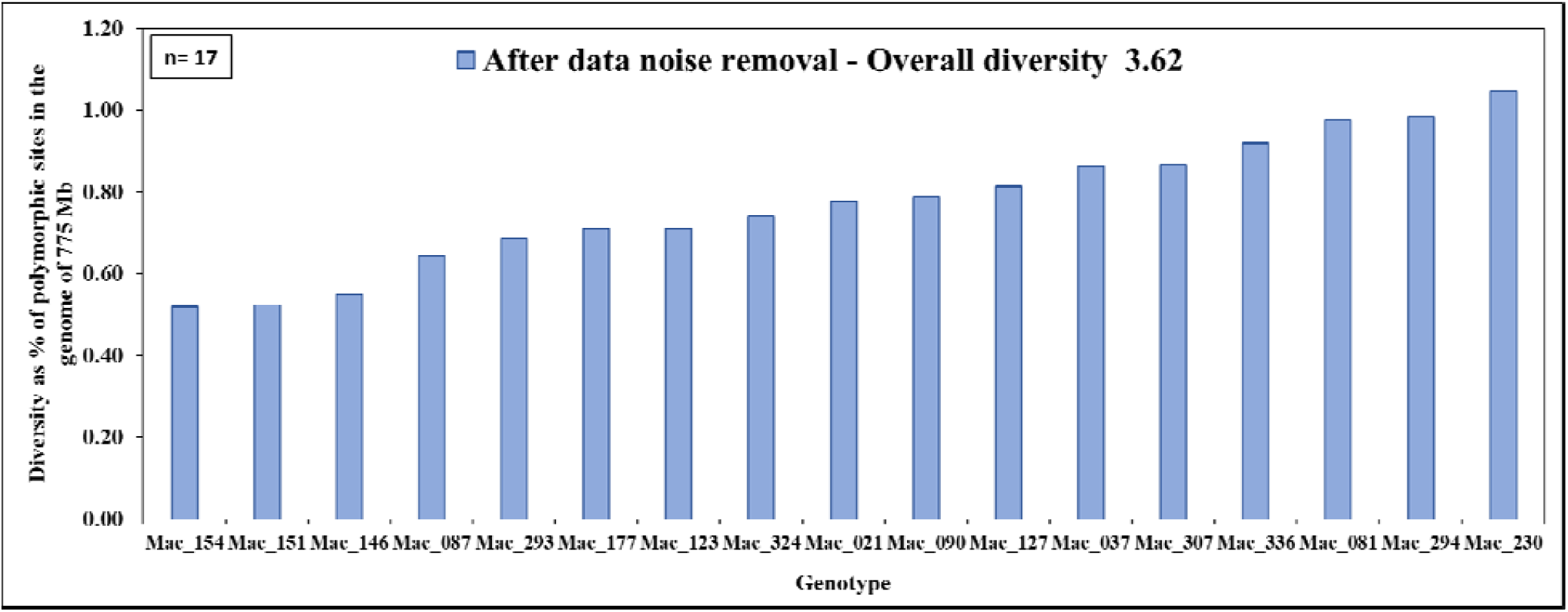
Diversity within Heritage cultivars. Diversity was calculated as a percentage of polymorphic sites in the genome of *M. integrifolia* (775 Mb)

**Figure S. 19.**
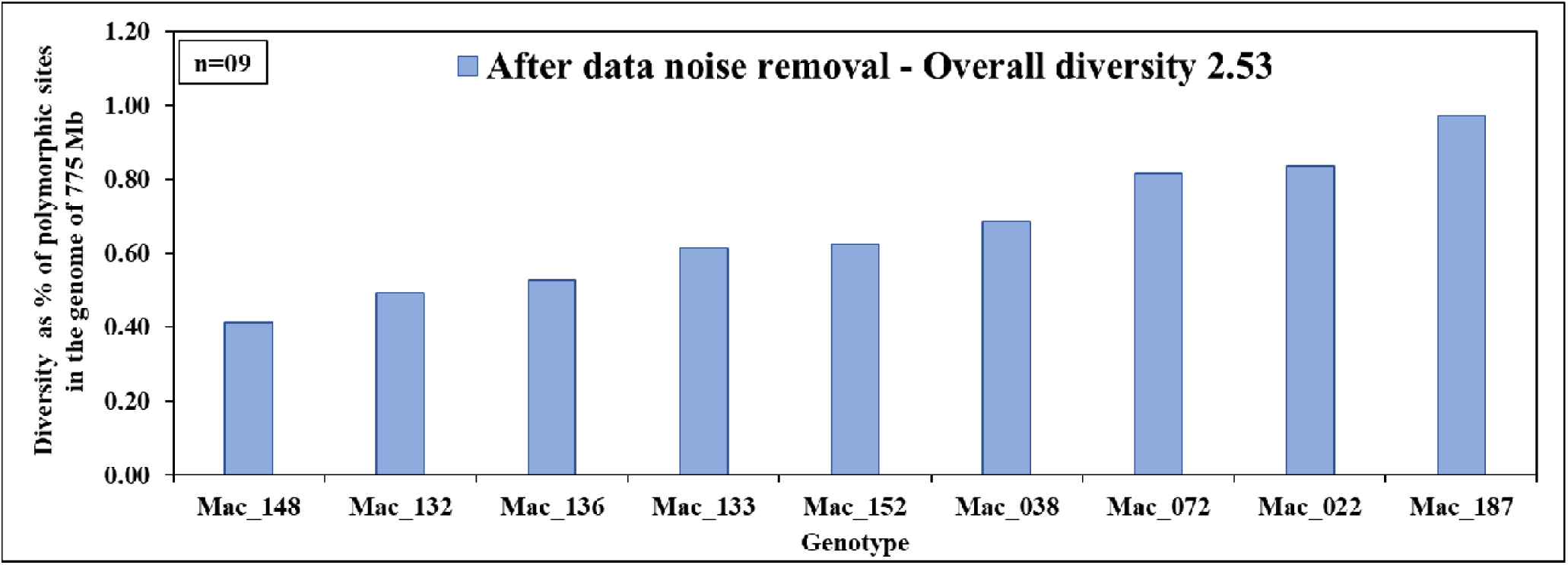
Diversity within cultivars bred by Norm Greber. Diversity was calculated as a percentage of polymorphic sites in the genome of *M. integrifolia* (775 Mb)

**Figure S. 20.**
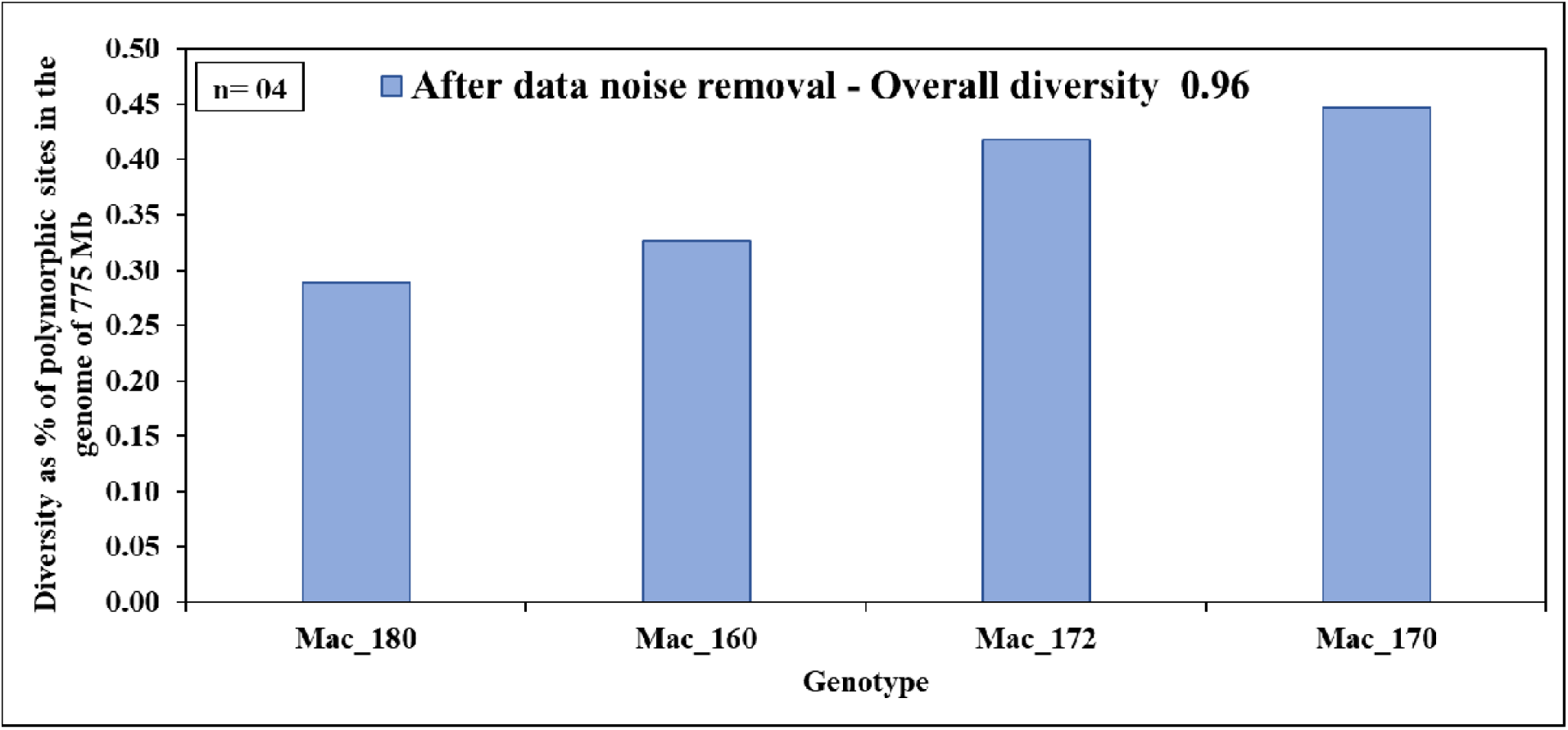
Diversity within cultivars bred by Backer. Diversity was calculated as a percentage of polymorphic sites in the genome of *M. integrifolia* (775 Mb)

**Figure S. 21.**
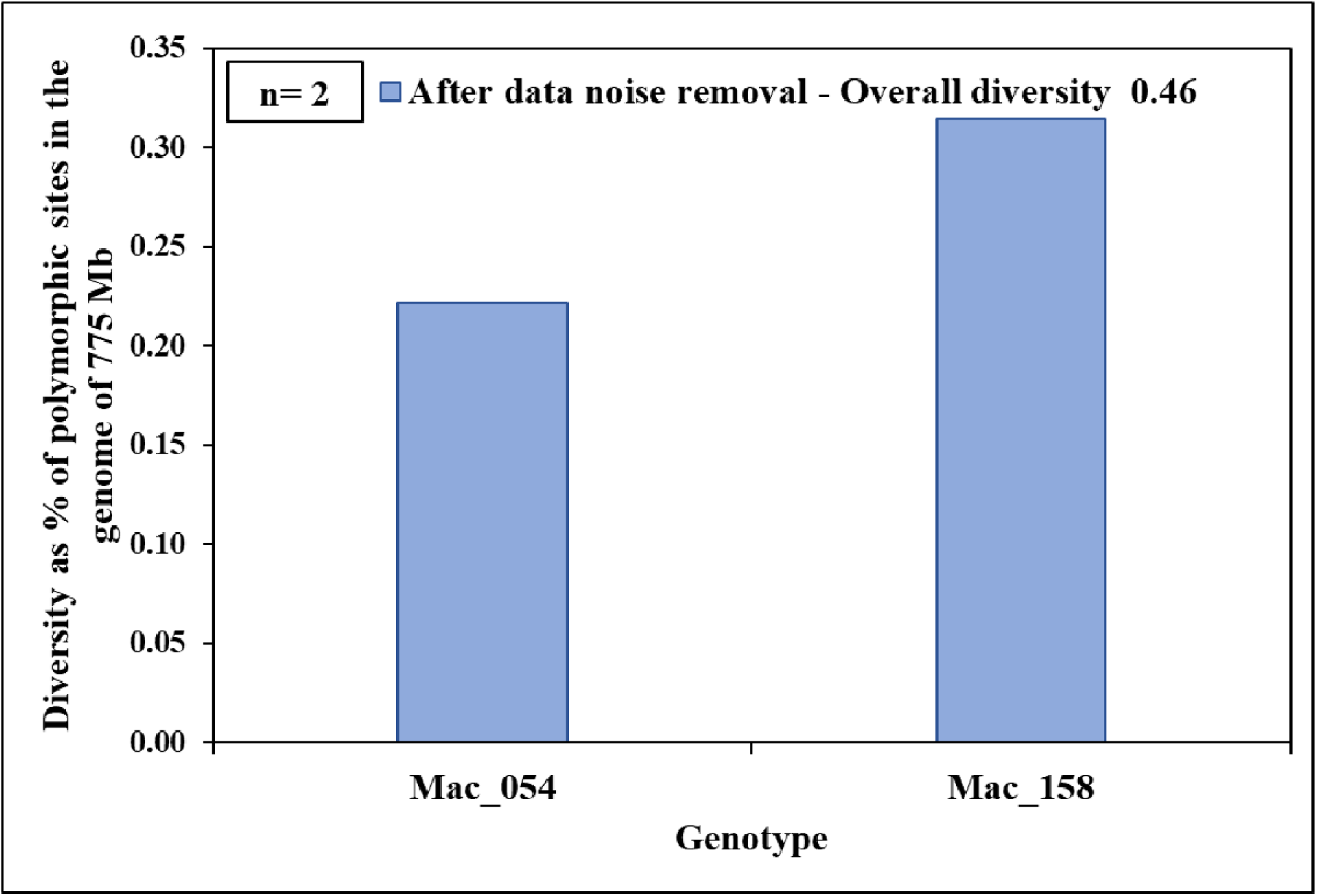
Diversity within cultivars bred by Malawi. Diversity was calculated as a percentage of polymorphic sites in the genome of *M. integrifolia* (775 Mb)

**Figure S. 22.**
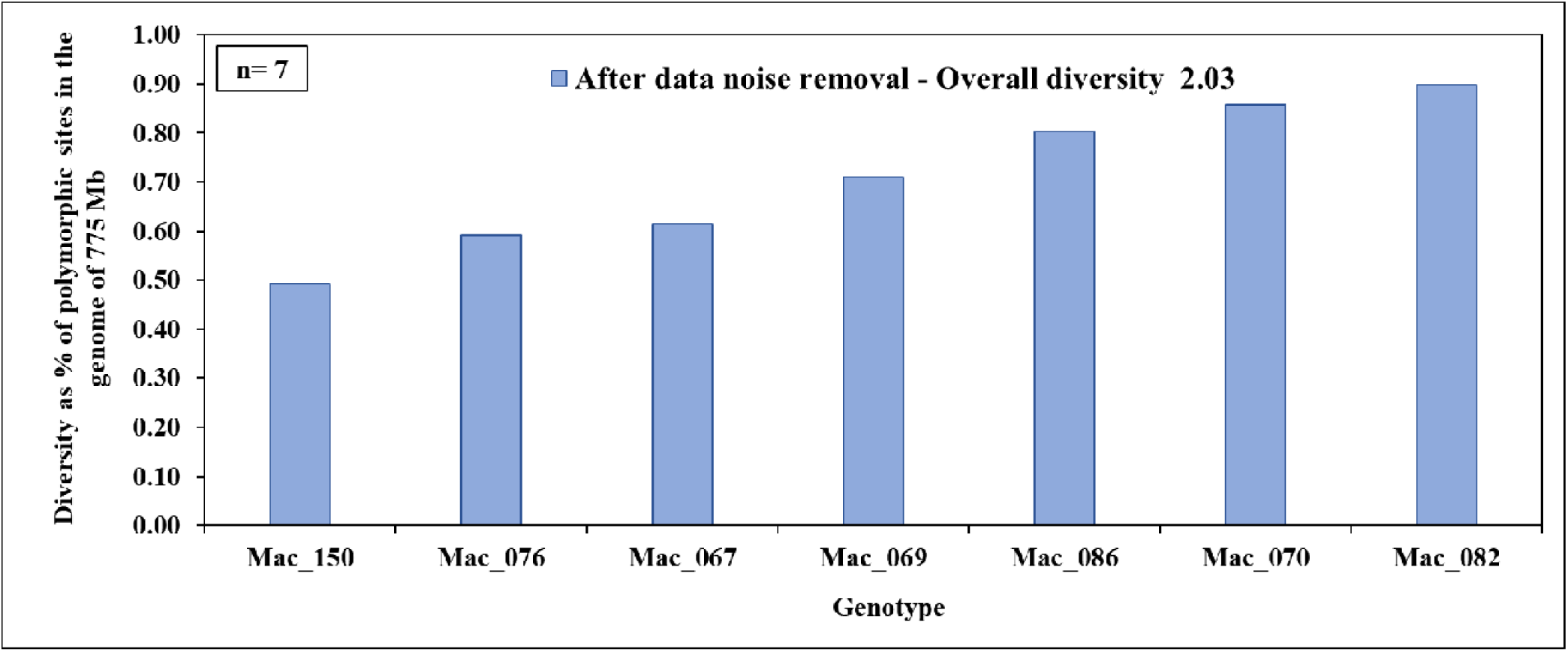
Diversity within cultivars bred by Ian McConachie. Diversity was calculated as a percentage of polymorphic sites in the genome of *M. integrifolia* (775 Mb)

**Figure S. 23.**
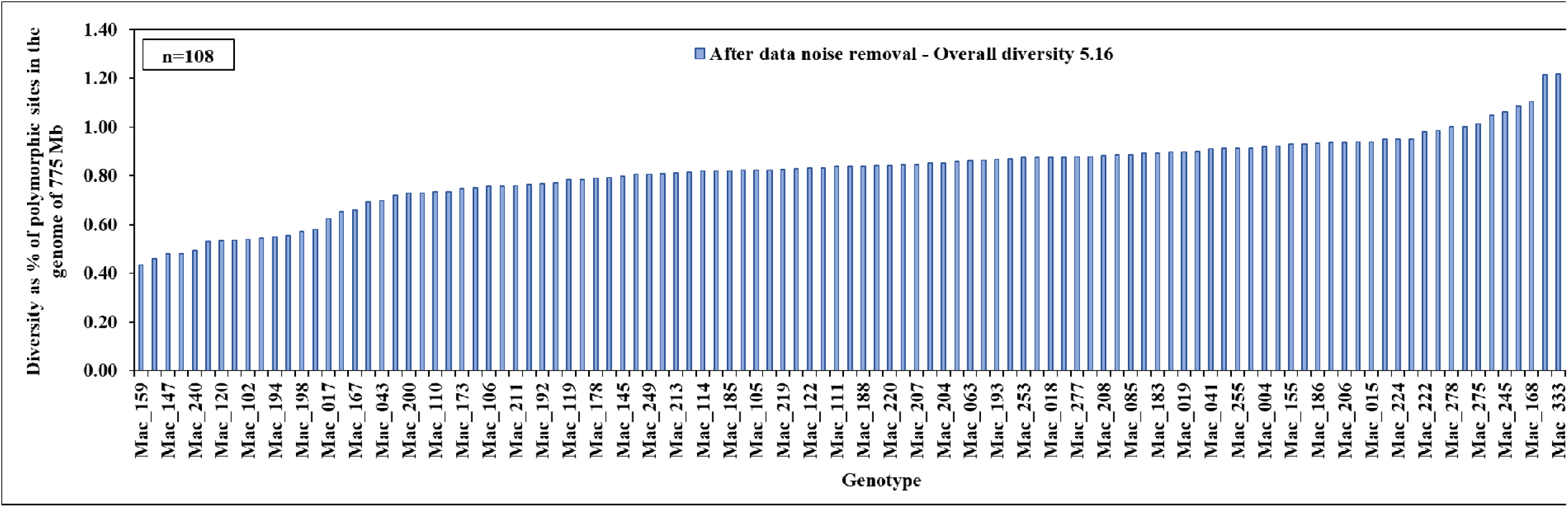
Diversity within Australian National Macadamia Breeding Program Accessions (NMBPA). Diversity was calculated as a percentage of polymorphic sites in the genome of *M. integrifolia* (775 Mb)

**Figure S. 24.**
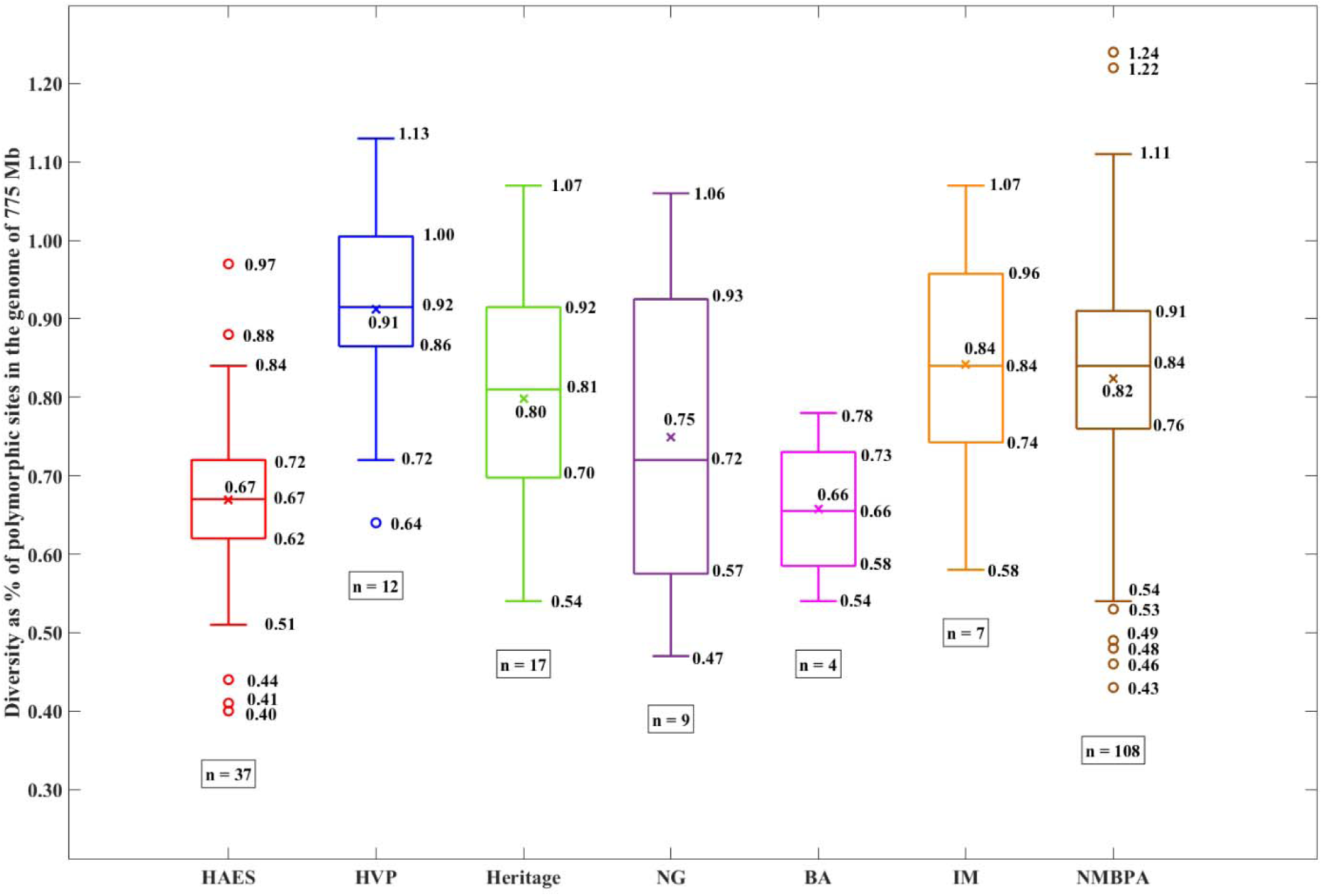
Comparison of diversity in different breeding programs. Diversity was calculated as a percentage of polymorphic sites in the genome of *M. integrifolia* (775 Mb). HAES: Hawaiian Agricultural Experiment Station cultivars. HVP: Hidden Valley Plantation cultivars. Heritage: Heritage cultivars. NG: Cultivars bred by Norm Greber. BA: Cultivars bred by Backer. IM: Cultivars bred by Ian McConachie. NMBPA: Australian National Macadamia Breeding Program Accessions.

**Figure S. 25.**
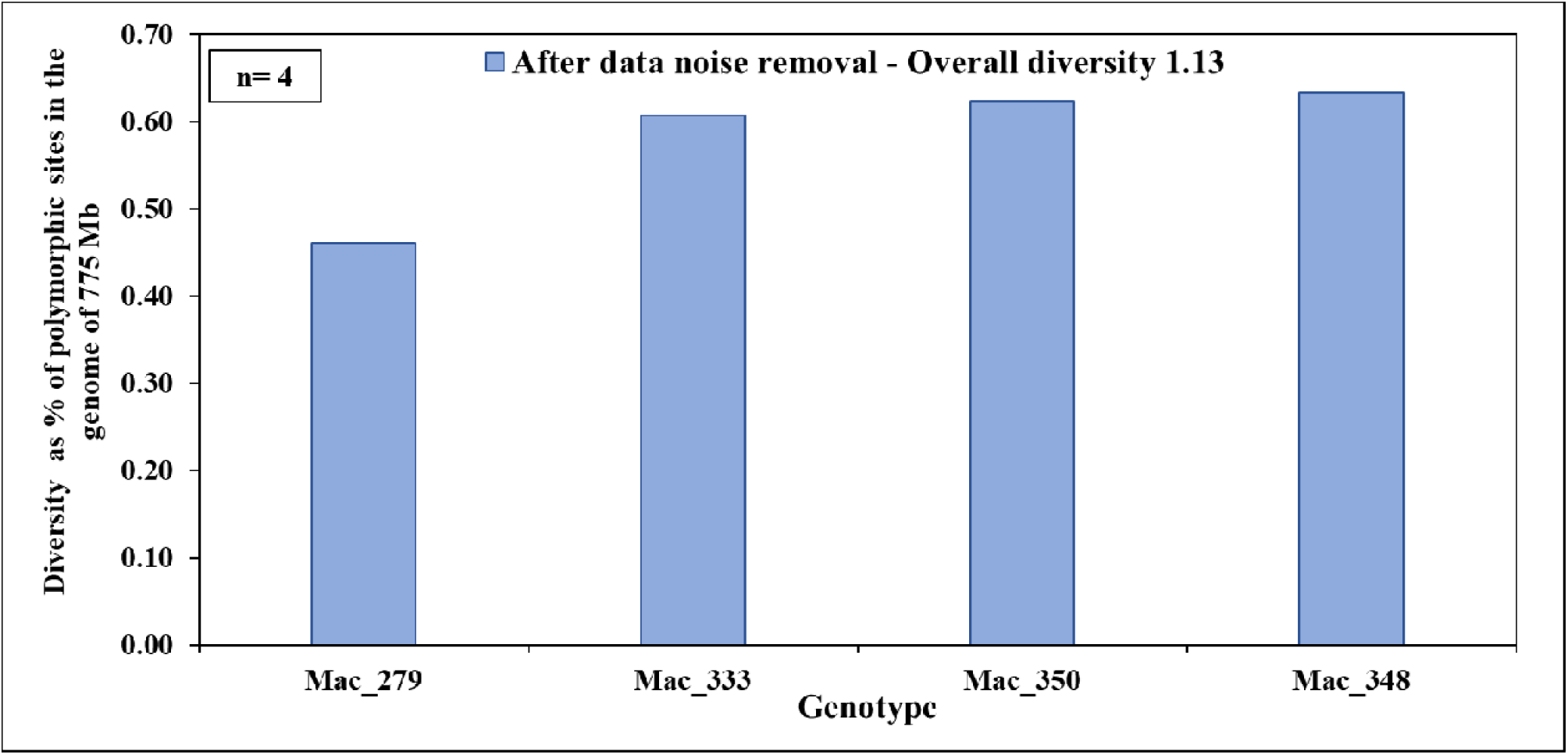
Diversity within genotypes grouped in clade I in nuclear phylogenetic analysis performed in our previous publication. Diversity was calculated as a percentage of polymorphic sites in the genome of *M. integrifolia* (775 Mb)

**Figure S. 26.**
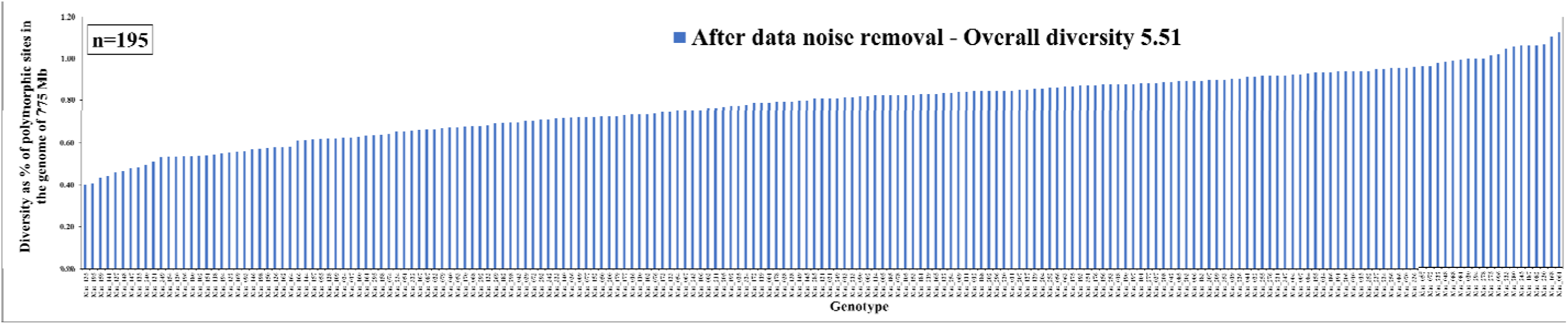
Diversity within genotypes grouped in clade II in nuclear phylogenetic analysis performed in our previous publication. Diversity was calculated as a percentage of polymorphic sites in the genome of *M. integrifolia* (775 Mb)

**Figure S. 27.**
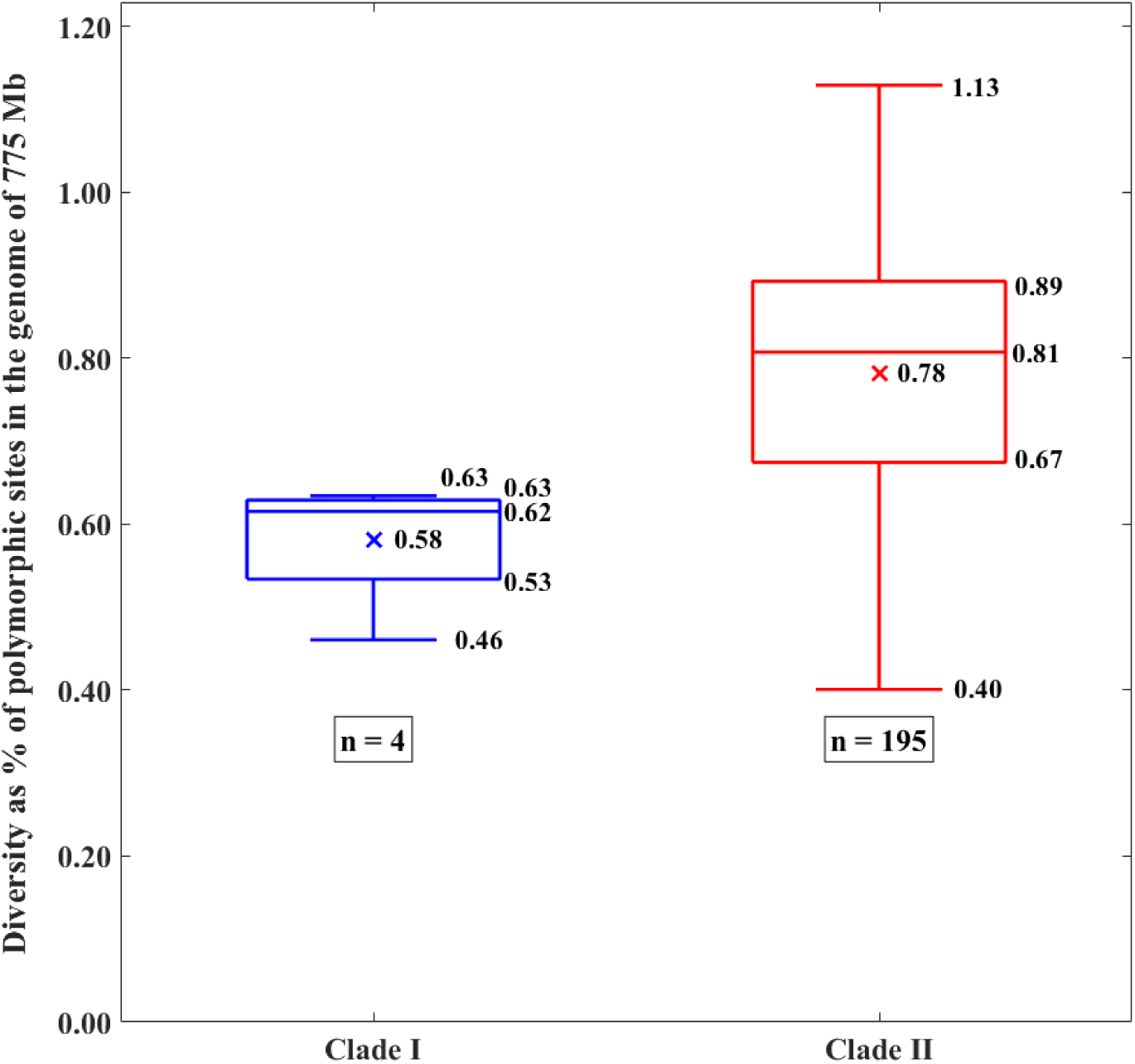
Comparison of diversity in clade I and clade II in nuclear phylogenetic analysis performed in our previous publication. Diversity was calculated as a percentage of polymorphic sites in the genome of *M. integrifolia* (775 Mb)

**Figure S. 28.**
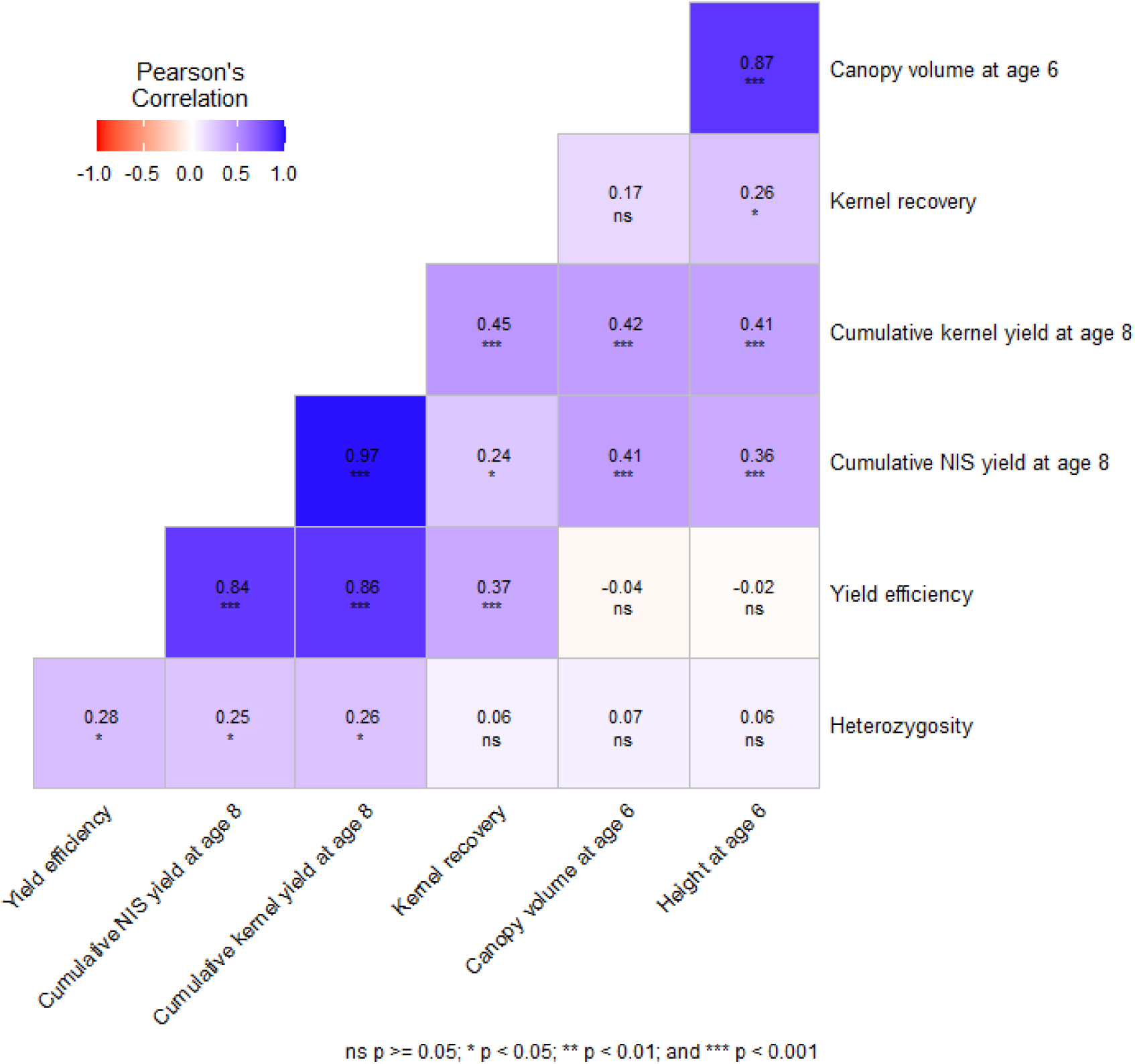
Correlation analysis results.

**Figure S. 29.**
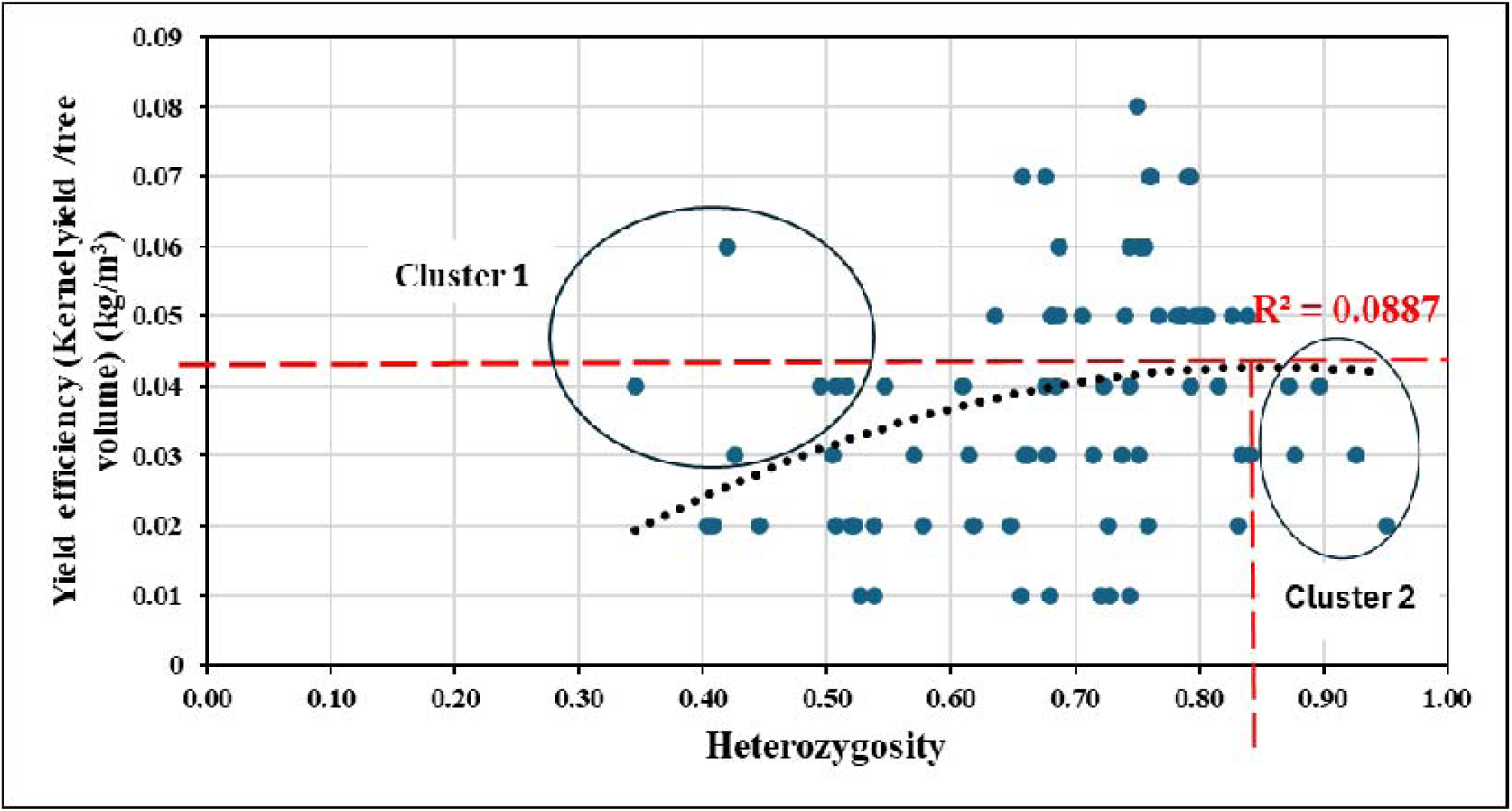
Relationship between the heterozygosity and yield efficiency (Kernel yield/tree volume).

**Figure S. 30.**
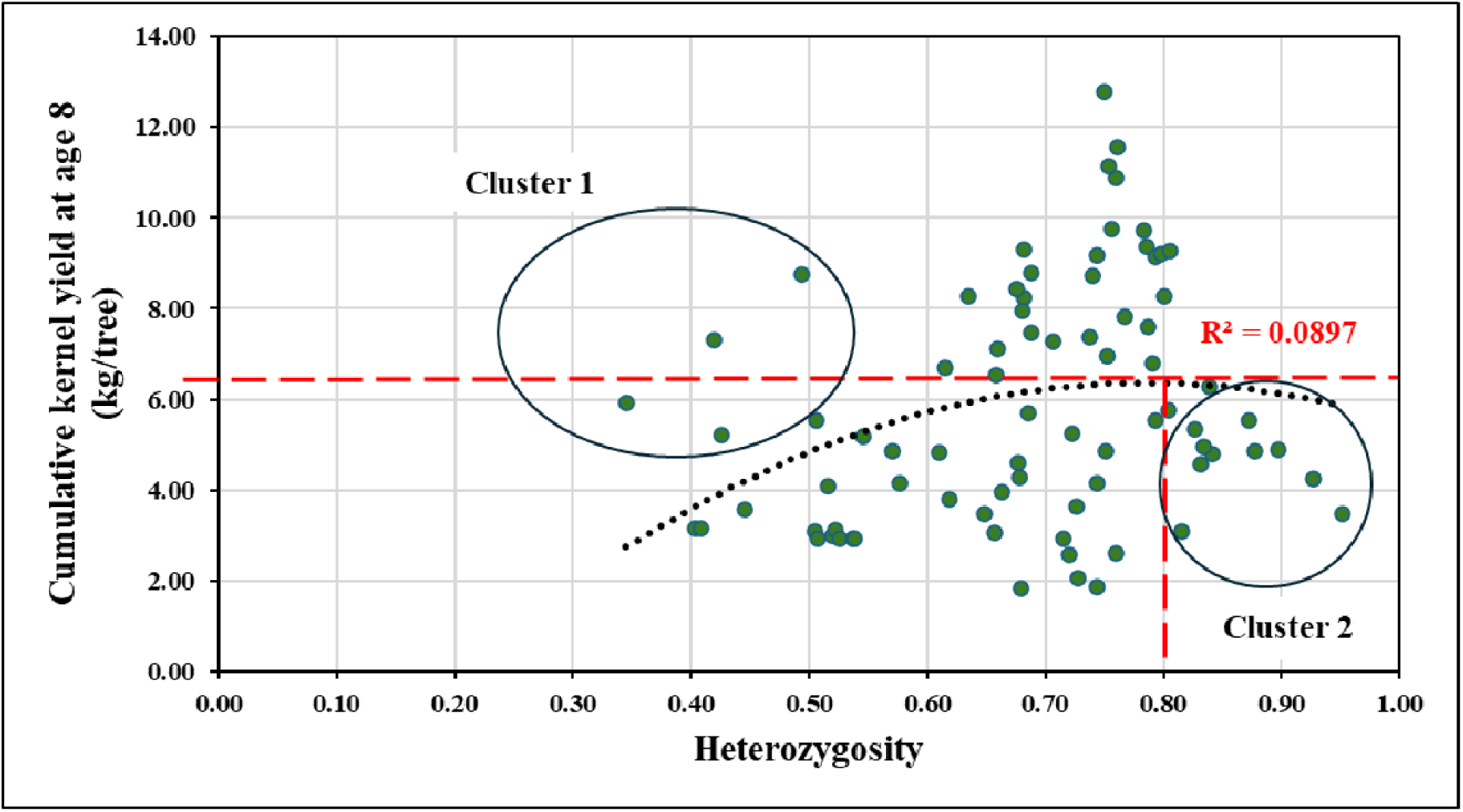
Relationship between the heterozygosity and cumulative kernel yield at age 8.

**Figure S. 31.**
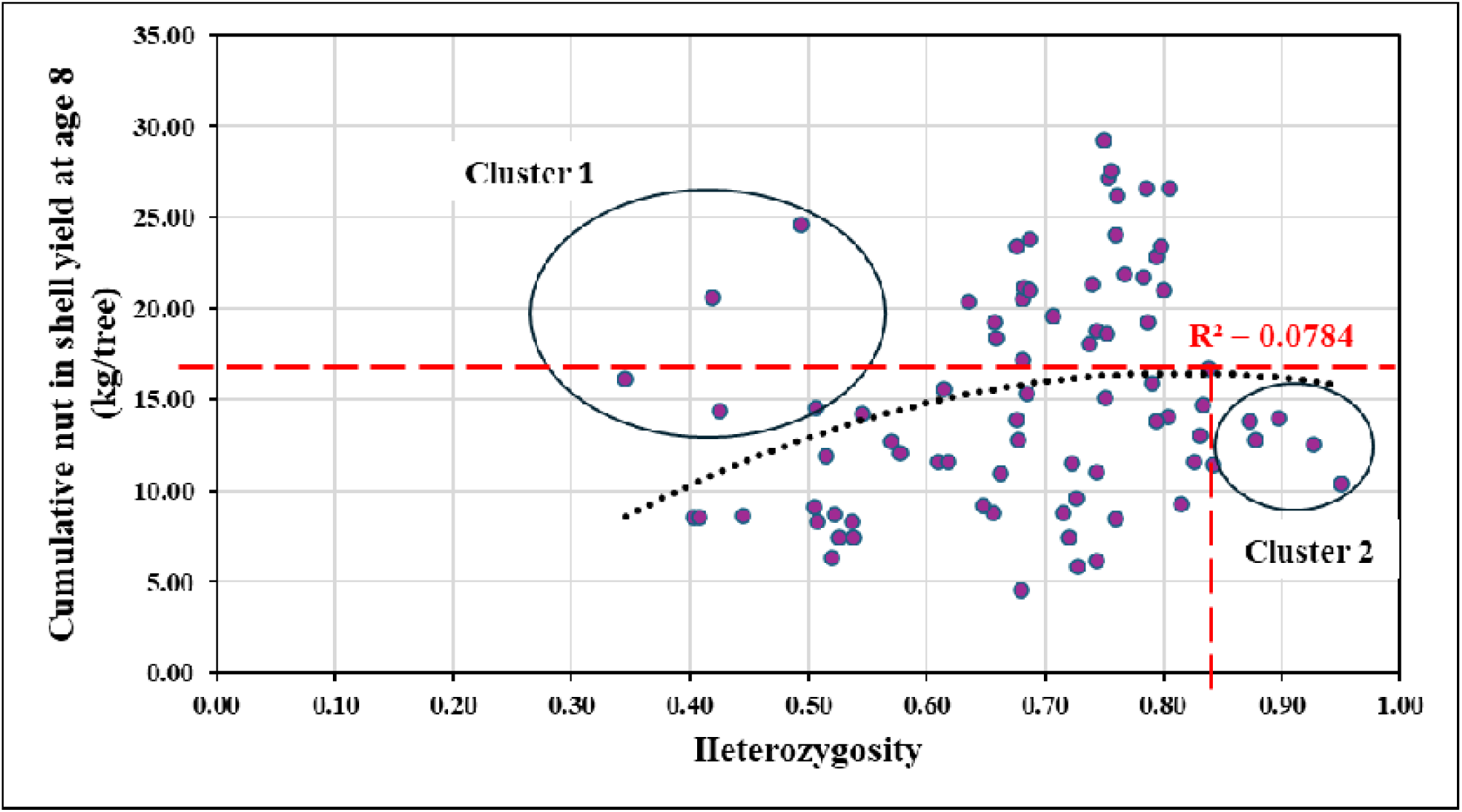
Relationship between the heterozygosity and cumulative nut in shell (NIS) yield at age 8.

**Figure S. 32.**
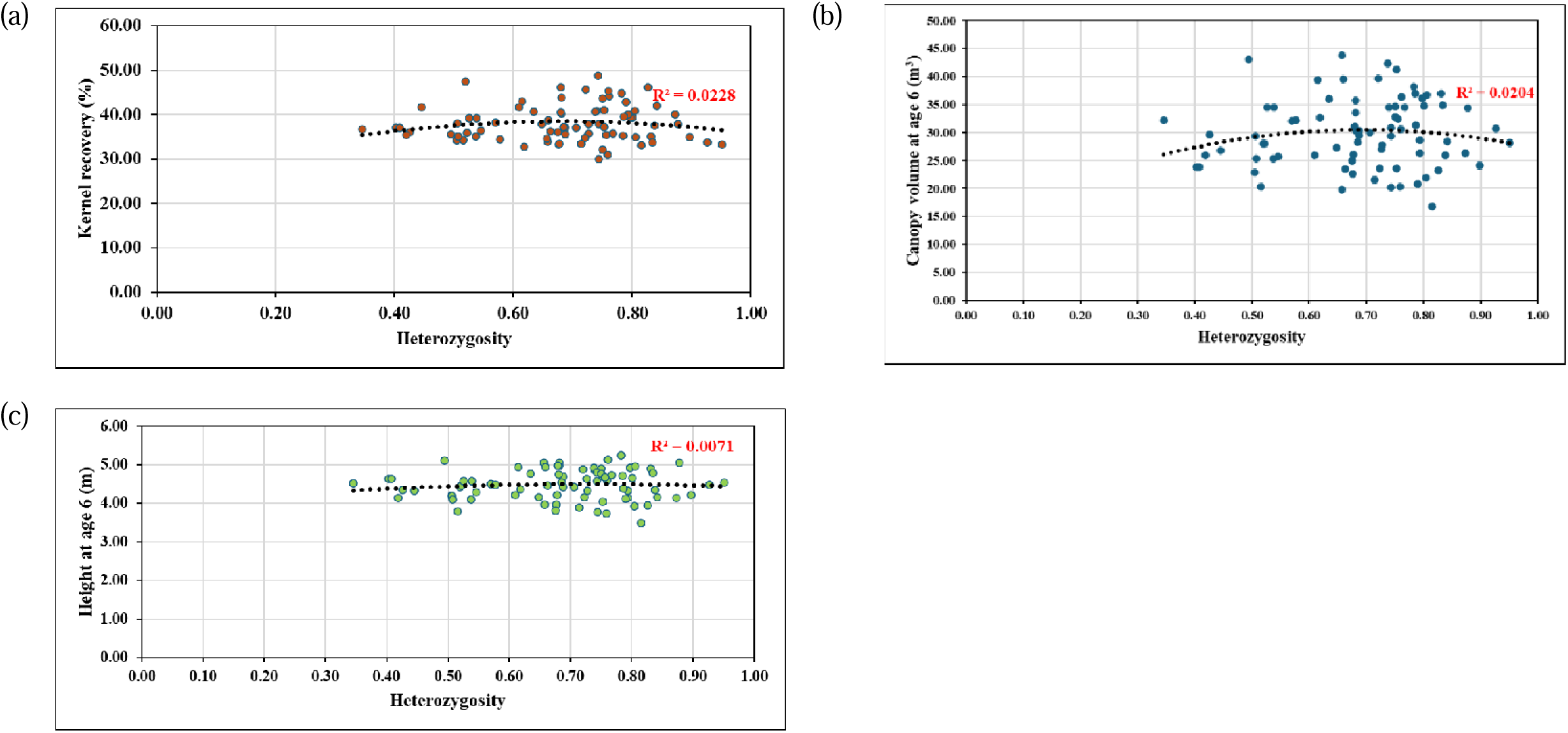
Relationship between heterozygosity and phenotypic traits. (a) Relationship between the heterozygosity and kernel recovery. (b) Relationship between the heterozygosity and canopy volume at age 6. (c) Relationship between the heterozygosity and tree height at age 6.

**Figure S. 33.**
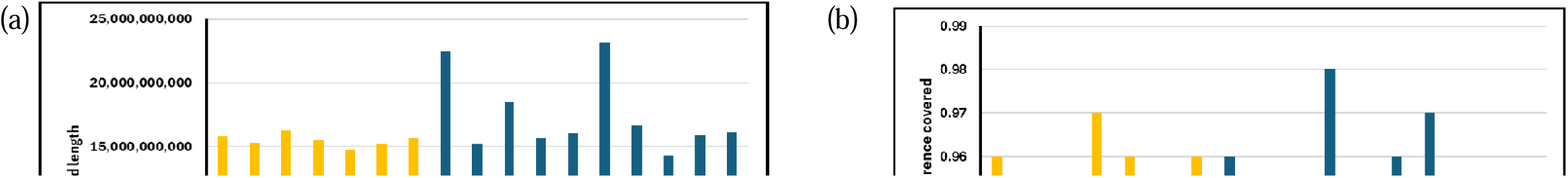
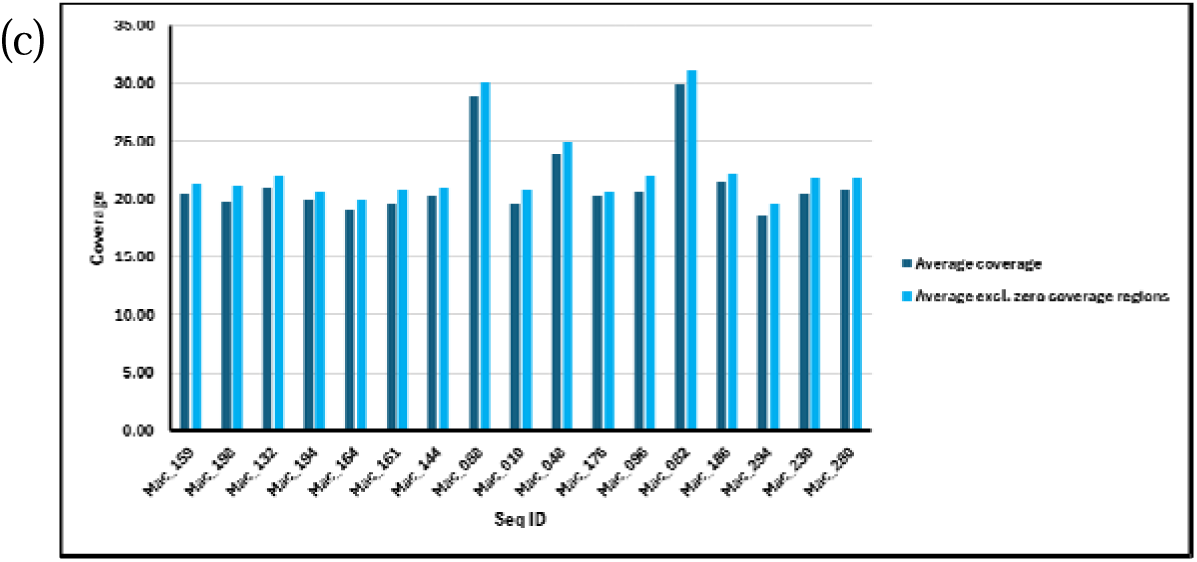
Coverage statistics. Mac_159, Mac_198, Mac_132, Mac_194, Mac_164, Mac_161 and Mac_144 are genotypes clustered in cluster 1 in Figure 6.11, 6.12 and 6.13. Mac_088, Mac_010, Mac_048, Mac_176, Mac_096, Mac_082, Mac_186, Mac_294, Mac_230 and Mac_280 are genotypes clustered in cluster 2 in Figure S.29, S. 30 and S. 31. (a) Graph shows the total read length between the genotypes in cluster 1 and 2. (b) Graph shows fraction of reference covered genotypes in cluster 1 and 2. 1/1 indicates the highest coverage. In (a) and (b) yellow indicates genotypes in cluster 1 and blue indicates genotypes in cluster 2. (c) Graph shows average coverage and average coverage excluding zero coverage region. Dark blue represents average coverage and light blue represents average coverage excluding zero coverage region

## List of References

1. Caliskan M: Genetic diversity in plants: BoD–Books on Demand; 2012.

2. El-Esawi MA: Introductory chapter: assessment and conservation of genetic diversity in plant species. In: Genetic Diversity in Plant Species-Characterization and Conservation. IntechOpen; 2019.

3. Yu Z, Fredua-Agyeman R, Hwang S-F, Strelkov SE: Molecular genetic diversity and population structure analyses of rutabaga accessions from Nordic countries as revealed by single nucleotide polymorphism markers. BMC genomics 2021, 22(1):442.

4. Xiong Y, Zhao Y, He Y, Zhao L, Hu H, Barrett RL, Chen Z, Lu L: Current progress and future prospects for understanding genetic diversity of seed plants in China. Biological Diversity 2024, 1(1):13–21.

5. Chung MY, Merilä J, Li J, Mao K, López-Pujol J, Tsumura Y, Chung MG: Neutral and adaptive genetic diversity in plants: An overview. Frontiers in Ecology and Evolution 2023, 11:1116814.

6. Bhandari H, Bhanu AN, Srivastava K, Singh M, Shreya HA: Assessment of genetic diversity in crop plants-an overview. Adv Plants Agric Res 2017, 7(3):279–286.

7. Pobke K: Draft recovery plan for 23 threatened flora taxa on Eyre Peninsula, South Australia 2007-2012. Department for Environment and Heritage, South Australia 2007.

8. Anderson JT, Willis JH, Mitchell-Olds T: Evolutionary genetics of plant adaptation. Trends in Genetics 2011, 27(7):258–266.

9. Topp BL, Nock CJ, Hardner CM, Alam M, O’Connor KM: Macadamia (Macadamia spp.) Breeding. In: Advances in Plant Breeding Strategies: Nut and Beverage Crops. 2019: 221–251.

10. Gross CL: Flora of Australia Elaeagnaceae Proteaceae 1. 1995:419–425.

11. Walton HMWaDA: Macadamia (Macadamia integrifolia, Macadamia tetraphylla and hybrids). In: Postharvest Biology and Technology of Tropical and Subtropical Fruits: Cocona to Mango. Edited by Yahia EM: Woodhead Publishing Limited.; 2011: 450–474.

12. Dahler JM, C. McConchie, and C. G. N. Turnbull: Quantification of cyanogenic glycosides in seedlings of three Macadamia (Proteaceae) species. Australian Journal of Botany 1995, 43(6):619–628.

13. Howlett BG, Read SF, Alavi M, Cutting BT, Nelson WR, Goodwin RM, Cross S, Thorp TG, Pattemore DE: Cross-pollination enhances macadamia yields, even with branch-level resource limitation. HortScience 2019, 54(4):609–615.

14. Jeong I-S, Yoon U-H, Lee G-S, Ji H-S, Lee H-J, Han C-D, Hahn J-H, An G, Kim T-H: SNP-based analysis of genetic diversity in anther-derived rice by whole genome sequencing. Rice 2013, 6:1–12.

15. Jiao F, Li T, Gao Y, Sun C, Wu X, Song Z, Chen X, Li Y, Gao L: Whole genome re-sequencing to reveal genetic diversity and to develop snp array in common tobacco. Biologia 2025:1–10.

16. Würschum T, Langer SM, Longin CFH, Korzun V, Akhunov E, Ebmeyer E, Schachschneider R, Schacht J, Kazman E, Reif JC: Population structure, genetic diversity and linkage disequilibrium in elite winter wheat assessed with SNP and SSR markers. Theoretical and Applied Genetics 2013, 126:1477–1486.

17. Cui J, Yang Y, Luo S, Wang L, Huang R, Wen Q, Han X, Miao N, Cheng J, Liu Z: Whole-genome sequencing provides insights into the genetic diversity and domestication of bitter gourd (Momordica spp.). Horticulture research 2020, 7.

18. Qi J, Liu X, Shen D, Miao H, Xie B, Li X, Zeng P, Wang S, Shang Y, Gu X: A genomic variation map provides insights into the genetic basis of cucumber domestication and diversity. Nature genetics 2013, 45(12):1510–1515.

19. Wei X, Shen F, Zhang Q, Liu N, Zhang Y, Xu M, Liu S, Zhang Y, Ma X, Liu W: Genetic diversity analysis of Chinese plum (Prunus salicina L.) based on whole-genome resequencing. Tree Genetics & Genomes 2021, 17(3):26.

20. McKenna A, Hanna M, Banks E, Sivachenko A, Cibulskis K, Kernytsky A, Garimella K, Altshuler D, Gabriel S, Daly M: The Genome Analysis Toolkit: a MapReduce framework for analyzing next-generation DNA sequencing data. Genome research 2010, 20(9):1297–1303.

21. Li H, Handsaker B, Wysoker A, Fennell T, Ruan J, Homer N, Marth G, Abecasis G, Durbin R, Subgroup GPDP: The sequence alignment/map format and SAMtools. bioinformatics 2009, 25(16):2078–2079.

22. Li R, Yu C, Li Y, Lam T-W, Yiu S-M, Kristiansen K, Wang J: SOAP2: an improved ultrafast tool for short read alignment. Bioinformatics 2009, 25(15):1966–1967.

23. Peace CP: Genetic characterisation of macadamia with DNA markers. 2005.

24. Peace C, Hardner C, Brown A, O’Connor K, Vithanage V, Turnbull C, Carroll B: The diversity and origins of macadamia cultivars. WANATCA Yearbook 2002, 26:19–25.

25. Alam M, Neal J, O’Connor K, Kilian A, Topp B: Ultra-high-throughput DArTseq-based silicoDArT and SNP markers for genomic studies in macadamia. PLoS One 2018, 13(8):e0203465.

26. O’Connor K, Kilian A, Hayes B, Hardner C, Nock C, Baten A, Alam M, Topp B: Population structure, genetic diversity and linkage disequilibrium in a macadamia breeding population using SNP and silicoDArT markers. Tree Genetics & Genomes 2019, 15(2):1–16.

27. Mai T, Alam M, Hardner C, Henry R, Topp B: Genetic Structure of Wild Germplasm of Macadamia: Species Assignment, Diversity and Phylogeographic Relationships. Plants (Basel) 2020, 9(6):714.

28. Ranketse M, Hefer CA, Pierneef R, Fourie G, Myburg AA: Genetic diversity and population structure analysis reveals the unique genetic composition of South African selected macadamia accessions. Tree Genetics & Genomes 2022, 18(2):15.

29. Nock C: Genetic diversity and population structure of wild and domesticated macadamia. In.; 2022.

30. Sharma P, Murigneux V, Haimovitz J, Nock CJ, Tian W, Kharabian Masouleh A, Topp B, Alam M, Furtado A, Henry RJ: The genome of the endangered Macadamia jansenii displays little diversity but represents an important genetic resource for plant breeding. Plant Direct 2021b, 5(12):e364.

31. Lin J, Zhang W, Zhang X, Ma X, Zhang S, Chen S, Wang Y, Jia H, Liao Z, Lin J et al: Signatures of selection in recently domesticated macadamia. Nat Commun 2022, 13(1):242.

32. Li Z, Wu C, Ma J, Geng J, Tao L, He X, Gong L: Genetic diversity analysis of macadamia germplasm in China based on whole-genome resequencing. Tree Genetics & Genomes 2024, 20(3):14.

33. Schmidt TL, Jasper ME, Weeks AR, Hoffmann AA: Unbiased population heterozygosity estimates from genome-wide sequence data. Methods in Ecology and Evolution 2021, 12(10):1888–1898.

34. Chen M, Song X, Wu S, Yu A, Wei X, Qiu J, Pei D: Genomic insights into genome-wide heterozygosity and its impact on walnut adaptive evolution and improvement. Molecular Breeding 2025, 45(6):1–19.

35. Aradhya MK, Yee LK, Zee FT, Manshardt RM: Genetic variability in Macadamia. Genetic Resources and Crop Evolution 1998, 45(1):19–32.

36. Peace CP, Vithanage V, Turnbull CG, Carroll BJ: A genetic map of macadamia based on randomly amplified DNA fingerprinting (RAF) markers. Euphytica 2003, 134(1):17–26.

37. Nock CJ, Elphinstone MS, Ablett G, Kawamata A, Hancock W, Hardner CM, King GJ: Whole genome shotgun sequences for microsatellite discovery and application in cultivated and wild Macadamia (Proteaceae). Appl Plant Sci 2014b, 2(4):1300089.

38. Nock CJ, Baten A, Mauleon R, Langdon KS, Topp B, Hardner C, Furtado A, Henry RJ, King GJ: Chromosome-Scale Assembly and Annotation of the Macadamia Genome (Macadamia integrifolia HAES 741). G3 (Bethesda) 2020, 10(10):3497–3504.

39. Spain CS, Lowe AJ: Genetic consequences of subtropical rainforest fragmentation on Macadamia tetraphylla (Proteaceae). Silvae Genetica 2011, 60(1-6):241–249.

40. Schmidt AL, Scott L, Lowe AJ: Isolation and characterization of microsatellite loci from Macadamia. Molecular Ecology Notes 2006, 6(4):1060–1063.

41. Langdon KS: Unravelling the genetics of macadamia: integration of linkage and genome maps. Southern Cross University; 2020.

42. Hardner C, Pisanu P, Boyton S: National Macadamia Germplasm Conservation Program. In.: CSIRO Plant Industry, Queensland Bioscience Precinct; 2004.

43. Furtado A: DNA extraction from vegetative tissue for next-generation sequencing. Methods Mol Biol 2014, 1099:1–5.

44. Sharma P, Masouleh AK, Constantin L, Topp B, Furtado A, Henry RJ: Genome sequences to support conservation and breeding of Macadamia. Tropical Plants 2024, 3(1).

45. Furtado A, Gunther T, Henry R: A Standard Protocol for Plant Variety Identification Based Upon Whole Genome Sequencing. 2025.

46. Topp B: Macadamia Second Generation Breeding and Conservation. In.; 2019.

47. Topp B, Hardner C, Neal J, Kelly A, Russell D, McConchie C, O’Hare P: Overview of the Australian macadamia industry breeding program. In: XXIX International Horticultural Congress on Horticulture: Sustaining Lives, Livelihoods and Landscapes (IHC2014): 1127: 2014. 45–50.

48. Al-Abdallat A, Karadsheh A, Hadadd N, Akash M, Ceccarelli S, Baum M, Hasan M, Jighly A, Abu Elenein J: Assessment of genetic diversity and yield performance in Jordanian barley (Hordeum vulgare L.) landraces grown under Rainfed conditions. BMC Plant biology 2017, 17:1–13.

49. Yirgu M, Kebede M, Feyissa T, Lakew B, Woldeyohannes AB, Fikere M: Single nucleotide polymorphism (SNP) markers for genetic diversity and population structure study in Ethiopian barley (Hordeum vulgare L.) germplasm. BMC Genomic Data 2023, 24(1):7.

50. Sun M: Effects of population size, mating system, and evolutionary origin on genetic diversity in Spiranthes sinensis and S. hongkongensis. Conservation Biology 1996, 10(3):785–795.

51. Zhang J, Zhang W, Ji F, Qiu J, Song X, Bu D, Pan G, Ma Q, Chen J, Huang R: A high-quality walnut genome assembly reveals extensive gene expression divergences after whole-genome duplication. Plant biotechnology journal 2020, 18(9):1848.

52. Yang Z, Ma W, Wang L, Yang X, Zhao T, Liang L, Wang G, Ma Q: Population genomics reveals demographic history and selection signatures of hazelnut (Corylus). Horticulture Research 2023, 10(5):uhad065.

53. Hu W, Ji C, Liang Z, Ye J, Ou W, Ding Z, Zhou G, Tie W, Yan Y, Yang J: Resequencing of 388 cassava accessions identifies valuable loci and selection for variation in heterozygosity. Genome biology 2021, 22:1–23.

54. Zeng L, Tu X-L, Dai H, Han F-M, Lu B-S, Wang M-S, Nanaei HA, Tajabadipour A, Mansouri M, Li X-L: Whole genomes and transcriptomes reveal adaptation and domestication of pistachio. Genome Biology 2019, 20:1–13.

55. Singh NK, Mahato AK, Jayaswal PK: The genome sequence and transcriptome studies in mango (Mangifera indica L.). The mango genome 2021:165–186.

56. Shirasawa K, Kosugi S, Sasaki K, Ghelfi A, Okazaki K, Toyoda A, Hirakawa H, Isobe S: Genome features of common vetch (Vicia sativa) in natural habitats. Plant Direct 2021, 5(10):e352.

57. Hardenbol P, Yu F, Belmont J, MacKenzie J, Bruckner C, Brundage T, Boudreau A, Chow S, Eberle J, Erbilgin A: Highly multiplexed molecular inversion probe genotyping: over 10,000 targeted SNPs genotyped in a single tube assay. Genome research 2005, 15(2):269–275.

58. Dai B, Guo H, Huang C, Zhang X, Lin Z: Genomic heterozygosity and hybrid breakdown in cotton (Gossypium): different traits, different effects. Bmc Genetics 2016, 17:1–11.

59. Savolainen O, Hedrick P: Heterozygosity and fitness: no association in Scots pine. Genetics 1995, 140(2):755–766.

60. Vranckx G, Jacquemyn H, Mergeay J, Cox K, Janssens P, Gielen BAS, Muys B, Honnay O: The effect of drought stress on heterozygosity–fitness correlations in pedunculate oak (Quercus robur). Annals of Botany 2014, 113(6):1057–1069.

61. López-Pujol J, Zhang F-M, Ge S: No correlation between heterozygosity and vegetative fitness in the narrow endemic and critically endangered Clematis acerifolia (Ranunculaceae). Biochemical genetics 2008, 46:433–445.

